# Sensory representation by non-equilibrium flows of neural activity at the mesoscale of cortex

**DOI:** 10.64898/2026.10.28.762488

**Authors:** Jochen Braun, Nick Lamb, Kingshuk Ghosh

## Abstract

We present a non-equilibrium theory of sensory perception and the mesoscale of sensory cortex. We propose that local neuron assemblies (cortical columns) toggle spontaneously between activity states (on and off) and that small populations of columns are coupled within and between areas of sensory cortex. The resulting network generates sudden, large-scale fluctuations and represents sensory inputs and perceptions by characteristic patterns of activity *flows*. The non-equilibrium dynamic is governed by the variational principle of maximal *path* entropy, or Maximum Caliber. Many other theories, by contrast, posit equilibrium dynamics and representation with activity *states*.

Our minimal model of sensory cortex describes populations of columns from two cortical areas and is fully constrained by empirical observations of ‘reversals’ in perceived appearance, which occur spontaneously for visual, auditory, and tactile stimulation. We formalize representation by activity *flows* with dynamic Hamiltonian functions. Surprisingly, effective representation requires small-number fluctuations of toggling columns, as larger numbers would be dominated by hysteresis. Finally, we detail predictions for the microscale of spiking neurons that follow from on-off toggling and non-equilibrium dynamics.

We conclude that the neural levels underlying perception are functionally organized in terms of activity *flows*. The inherent instability of non-equilibrium ensures active updating and renders all perceptions open to immediate revision. We conjecture that current theories of brain function (such as the Free Energy Principle) could be reformulated in terms of the variational principle of non-equilibrium, Maximum Caliber.

## Introduction

Like all living systems (1), the brain relies extensively on non-equilibrium processes to perform essential functions. At the *microscale* of individual neurons, irreversible flows contribute prominently to electrical excitability, active transport, synaptic transmission, and many other functions (2, 3). At the *macroscale* of whole-brain activity, neural dynamics exhibit all the hallmarks of non-equilibrium behaviour, such as unbalanced (4–6) or turbulent flows (7, 8). This large-scale irreversibility may also relate to function, as it correlates with levels of wakefulness (9, 10), cognitive engagement (11), and developmental maturity (12).

At the *mesoscale* of the sensory cortex, where local assemblies of neurons (‘cortical columns’) support visual, auditory, and tactile perception (13–15), the possibility of nonequilibrium activity has rarely been considered. At this scale, neuronal activity varies spontaneously and is low-dimensional (relative to the number of neurons), transitioning between a small repertoire of distinct states (16–18). More specifically, the spontanenous variability of activity is largely shared between nearby neurons (37–39) and local assemblies can transition spontaneously between distinct ‘on’ and ‘off’ states (19–21). Precise mathematical laws (mean-field equations) describe such ‘multistable attractor’ dynamics by assemblies of spiking neurons (22–26). Because unprompted state transitions are metabolically costly, it is natural to wonder about their functional relevance (27).

Here we present a non-equilibrium theory for the activity of sensory cortex at the mesoscale of toggling cortical columns. We suggest that small populations of columns are coupled through competitive, cooperative, and destabilizing interactions and collectively express sudden, large-scale fluctuations. Large fluctuations of mesoscale activity are known to accompany both stimulus-evoked (28–30) and spontaneous (31–33) changes in perception. More specifically, we propose that mesoscale activity involves unbalanced flows and non-equilibrium steady states (NESS). We show that large fluctuations of mesoscopic activity can be described by the variational principle of non-equilibrium known as Maximum Caliber (34, 35).

What sets our theory apart from other views is that sensory inputs and perceptions are represented jointly by characteristic patterns of activity *flows*, rather than by patterns of activity *states* (36). A pattern of flows is defined by particular distribution of path probabilities and can be formalized by a dynamical Hamiltonian function. Also, our theory suggests that the shared variability of cortical activity mentioned above (37–39) reflects degeneracy of toggling columns. Thus, we add to the growing body of work that places *trajectories* through state space at the center of neural computation and representation (40–43).

In our scenario, activity states remain far from equilibrium, and external sensory input shapes the ‘topologies of conductivity’ that steer activity flow in local cortical networks. In many other theories of sensory decision-making (26, 44, 45) and hierarchical inference (such as the Free Energy Principle) (46–48), the activity states of local networks perform noise-driven random walks, and sensory input shapes the ‘potential landscapes’ that turn random walks into gradient descents. Thus, cortical activity is governed by the variational principle of equilibrium (Maximum *State* Entropy) and evolves toward equilibrium states with balanced flows. We conjecture that these theories could be reformulated in terms of unbalanced flows and the variational principle of non-equilibrium (Maximum *Path* Entropy).

We illustrate our theory with small mesoscale models that were originally developed to comprehensively reproduce the rich dynamics of perceptual ‘reversals’ as experienced and reported by human observers (49). Sensory perception generally exhibits an intrinsic dynamic in that perceived appearances change spontaneously and repeatedly while sensory stimulation remains constant (50–53). These perceptual ‘reversals’ are endogenous (54, 55) and occur across a wide range of situations (56, 57) and sensory modalities (vision, audition, touch) (58–60) in mammals and birds (61–65). In primates, perceptual reversals are associated with activity transients at various levels of sensory cortex (66–68).

The statistics of perceptual reversals in human observers suggest that the underlying neural activity involves on–off toggling and operates far from equilibrium (i.e., far from saturation or depletion) (69, 70). This is what allows all aspects of our mesoscale models – number of columns, toggling rates, and interaction strengths – to be fully specified and constrained by this empirical data. Because reversal statistics are highly consistent across situations that engage different cortical areas (71–73), non-equilibrium toggling could be widespread in sensory cortex.

We report the dynamics of mesoscale models with two or three dominance alternatives under steady-state sensory input. We show that the dynamics constitute non-equilibrium steady states (NESS) formalized by dynamical Hamiltonian functions. Further, we show that smaller numbers of toggling columns (approximately two dozen per population) ensure sensitivity to sensory input. With larger numbers, the dynamics are trapped in limit cycles and lose their sensitivity to input. Finally, we show that the interaction strengths inferred from perception maximize path entropy and keep trajectories only weakly committed to any single dominance alternative. We conclude that these architectures are functionally organized for sensory decision-making.

Because our theory bridges neural scales from the macroscopic (perception) through the mesoscopic (toggling columns) to the microscopic (individual neurons), it makes quantitative predictions that can be tested neurophysiologically. One set of predictions concerns shared variability arising from the on–off toggling of local neuron assemblies with reciprocal interactions. Another set concerns arrow-of-time irreversibility (‘broken detailed balance’) arising from non-equilibrium trajectory flows (5, 74, 75) between different levels of the architecture, where interactions are non-reciprocal and mutually destabilizing.

## Results

We investigate mesoscale models of perceptual reversals between two or three alternatives (**Fig. 1**). Reversals between multiple alternatives are common in perception (76–79). Our models receive multiple sensory inputs and their intrinsic fluctuations almost always identify one perceptual alternative as ‘dominant’. While this identification can be correct and reflect actual differences between inputs (80), it can also be spurious, for example, when inputs are identical.

**Fig. 1.**
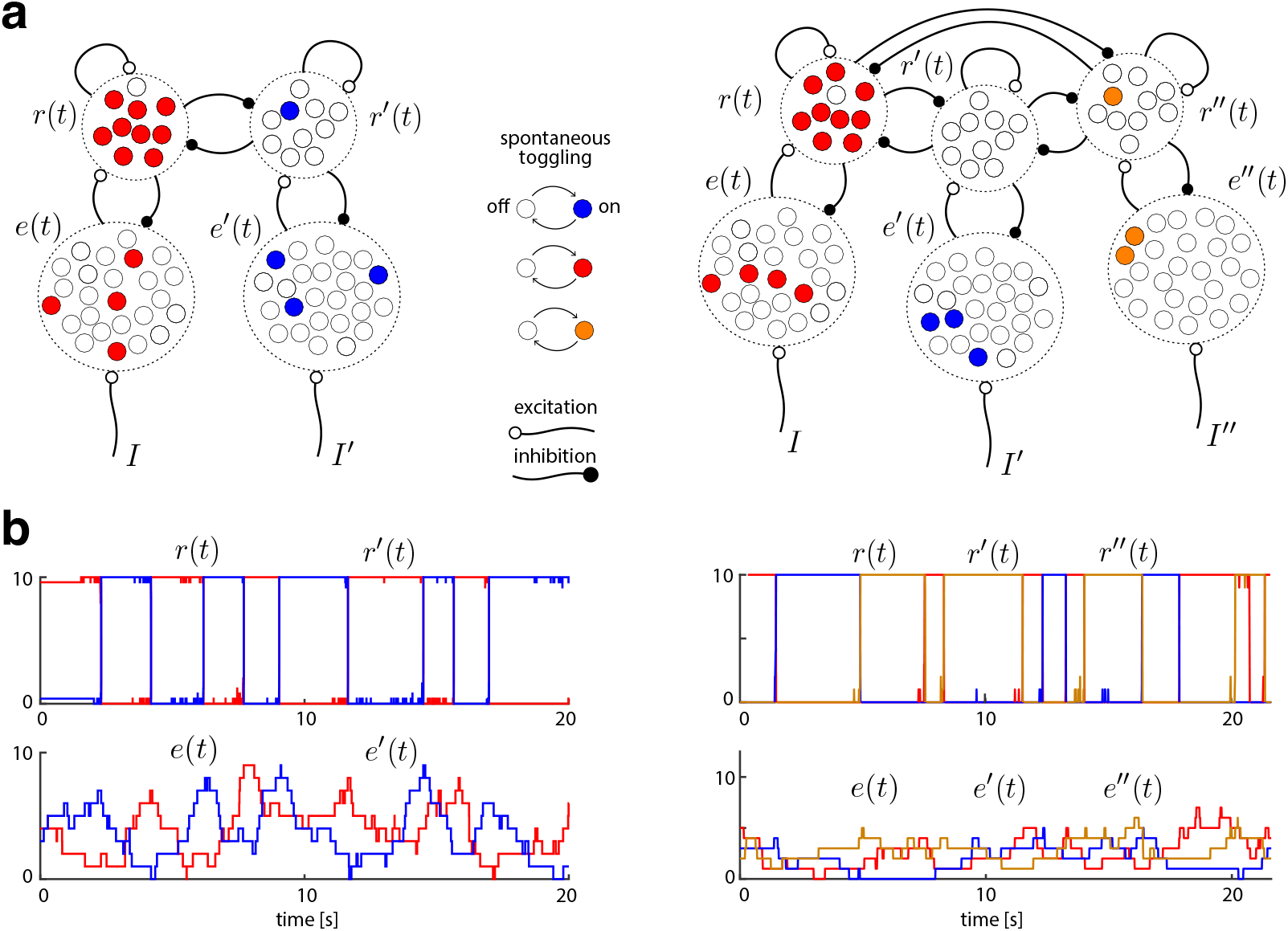
Neural architectures inferred from perceptual ‘reversals’ generate intrinsic fluctuations. Local assemblies of neurons or ‘columns’ (small circles) are bistable and toggle spontaneously and independently between ‘on’ and ‘off’ states. The collective activity of a population of columns (dashed circles) is a birth-death process, with external input to a population setting the toggling rates. Lower level ‘evidence’ populations, with activities *e, e*′, *e*″, receive (generally different) visual inputs, *I, I*′, *I*″, integrating sensory evidence over time. Upper level ‘dominance’ populations, with activities *r, r*′, *r*″, interact asymmetrically with lower level populations and competitively with each other (mutual inhibition balanced by self-excitation). As a result, upper-level populations almost always identify one particular input as ‘dominant’ (correctly or not). **(a)** Architectures with two (left) and three perceptual alternatives (right). **(b)** Representative stochastic trajectories generated by two- (left) and three-alternative architectures (right). Activity fluctuates spontaneously even though sensory input was equal and constant here, *I* = *I*′ = *I*″ = 0.5. Lower-level activities increase or decrease in individual steps (toggles), accumulating or discounting sensory evidence. Upper-level activities reverse their ‘dominance’ choices categorically (‘winner-take-all’).

When several bistable columns receive the same input, their combined activity becomes a birth-death process, which is driven towards saturation (all columns ‘on’) by excitatory input and towards depletion (all ‘off’) by inhibitory input. During such a stochastic accumulation, the combined activity acquires Fisher information about the average strength of the driving input. The rate of information uptake grows linearly with the number of columns (see further below).

Mesoscale models have two levels (upper and lower) and every perceptual alternative is represented by one birth-death population at each level (**Fig. 1a**). Upper-level activities represent ‘dominance’ and are mostly categorical (all-or-nothing), due to dynamic competition between alternatives (mutual inhibition balanced by self-excitation). Lower-level activities represent ‘evidence’ and are gradually increased by sensory input, but also decreased by the corresponding upper-level activity (**Fig. 1b**). This negative feedback ensures spontaneous reversals even when inputs remain unchanged.

As this model architecture was inferred from perceptual evidence, its neural substrate is not known with any certainty. However, the functional anatomy of primate visual cortex (81–83) suggests an initial hypothesis. We conjecture that one mesoscale circuit corresponds to a patch of cortex with the same ‘receptive field’ at one level of the visual cortical hierarchy, plus the directly afferent patches at the next lower level with smaller receptive fields (approximately O(10^3^) columns and 1 mm^2^ surface area). Presumably, the visual cortex as a whole (approximately 25.000 mm^2^ area) comprises some tens of thousands of replica circuits.

### Non-equilibrium steady state (NESS) dynamics

The non-equilibrium dynamics of mesoscale models become evident when trajectory flow is visualised in state space. Because the state space of the two-alternative model is four-dimensional (*e, e*′, *r, r*′), the average flow is best illustrated by short segments of average trajectories that start from selected initial locations and are projected into the two-dimensional subspace (*e − e*′, *r − r*′) (**Fig. 2a**) or the three-dimensional subspace (*e − e*′, *r − r*′, *e* + *e*′) (**Fig. 2b,c**). Crucially, the average flow in the steady state possesses a predominant direction and is therefore unbalanced, violating the definition of thermodynamic equilibrium (i.e., detailed balance is broken). In **Fig. 2a,b** this predominant direction is indicated by dotted loops.

**Fig. 2.**
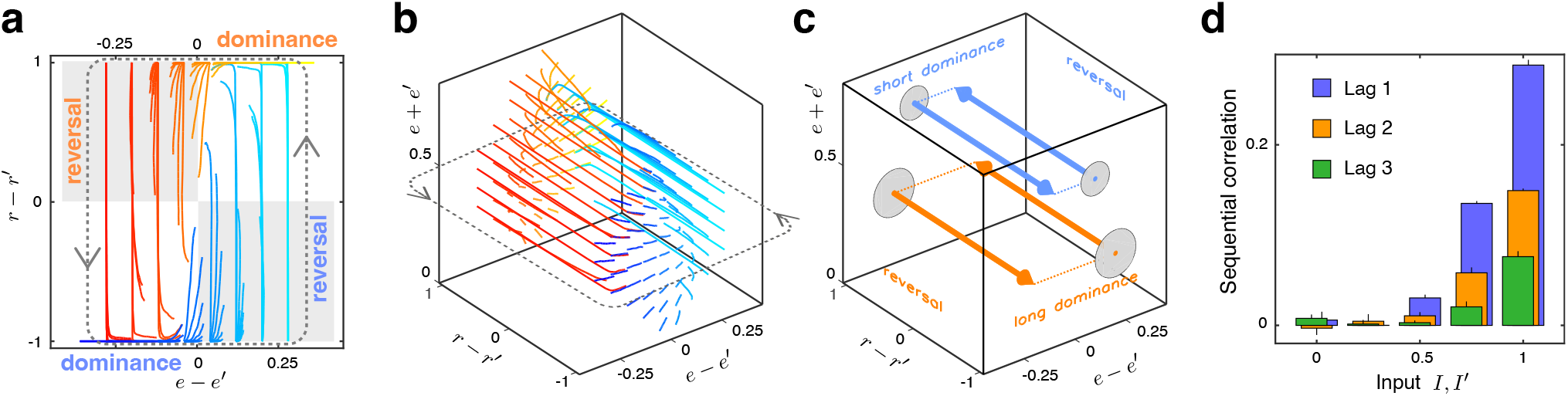
Non-equilibrium dynamics and sequential correlations. Average trajectory flow with dominance and reversal phases for two-alternative dynamics. For visualization, the 4D state space (*e, e*′, *r, r*′) is projected into 2D subspace (*e − e*′, *r − r*′) or 3D subspace (*e − e*′, *r − r*′, *e* + *e*′). Dotted loops indicate the predominant direction of flow. **(a)** Average trajectory flows of 0.2 s duration from 128 starting locations (the most frequently visited locations), distinguished by color. Trajectories proceed counter-clockwise, from yellow to red to blue to cyan and back to yellow. During ‘dominance phases’, the value of *r − r*′ remains mostly constant (near either *−*1 or +1) while the value of *e – e*′ changes. During ‘reversal phases’, the opposite is true. Reversals are triggered by ‘tension’ from opposite signs of *r − r*′ and *e − e*′ (grey areas). **(b)** Average trajectory flows in 3D space. Trajectories traverse more slowly (over shorter distances) during dominance phases and more rapidly (over longer distances) during reversal phases. Due to stochastic fluctuations, reversals may occur at different levels of *e* + *e*′. **(c)** At higher levels of *e* + *e*′, less ‘tension’ is needed for reversals, rendering dominance phases shorter (blue arrows and dashed lines), and *vice versa* for lower levels of *e* + *e*′ (orange arrows and dashed lines). **(d)** Due to slow random fluctuations in the level of *e* + *e*′, several successive dominance durations are correlated positively (lag 1, 2, and 3). Correlations increase with sensory input, in quantitative agreement with human perception (49).

Trajectory segments traverse different distances in state space, reflecting differences in speed. Shorter (slower) segments course within or near an (*e, e*′) plane at fixed values of (*r, r*′) (either *r − r*′ = 1 or *r − r*′ = *−*1), whereas longer (faster) segments traverse between the categorical values of (*r, r*′). The slow and fast segments correspond, respectively, to the dominance and reversal phases of the dynamics.

The alternation of dominance and reversal phases is sustained by the periodic build-up and release of internal tension. When one upper-level population dominates the other, inhibitory feedback drives the lower-level populations in the opposite direction, thereby generating a destabilising tension. Once the contradiction between *e − e*′ and *r − r*′ becomes sufficiently strong, the upper level is destabilised and dominance reverses, initiating a new cycle.

For the three-alternative dynamics the process is analogous, except that tension can be released by trajectories turning in either of two directions (corresponding to the two alternatives to the currently dominant state). Because the choice of direction is governed by non-equilibrium fluctuations, average trajectories are no longer informative. We return to this important point below.

Trajectories traverse different regions of state space, sometimes at lower and sometimes at higher levels of *e* + *e*′ (**Fig. 2c,d**). Slow stochastic fluctuations in the level of *e* + *e*′ introduce a history dependence that grows with sensory input: dominance periods of similar duration (either shorter or longer) tend to follow one another. This occurs because reversals are initiated at smaller differentials |*e − e*′| when *e* + *e*′ is larger, and vice versa. The model therefore predicts positive sequential correlations between successive dominance durations over several lags (lag-1, lag-2, etc.). Both the predicted sequential correlations and their dependence on sensory input have been confirmed quantitatively in human observers (49). This quantitative agreement in non-stationary aspects of the dynamics provides additional support for the mesoscale architecture.

### Sensory input evokes characteristic NESS flow

A very general approach to analysing non-equilibrium dynamics is to model the observed trajectories directly (35, 84, 85) by means of the variational principle of maximum path entropy, better known as Maximum Caliber (34, 86). Maximum Caliber is the dynamical counterpart of the maximum-entropy principle used for equilibrium systems. It assigns probabilities to entire paths by maximising path entropy subject to dynamical observables (constraints). Here we invert the procedure: we infer the constraints from the observed stochastic fluctuations of the system components. Typical constraints are the effective interactions between components; their strengths can be recovered from the stochastic trajectories. The end product is a dynamical Hamiltonian that depends on the instantaneous system state, encodes the effective interactions, and determines the probability of every possible state transition. Because it relies solely on observed correlations and on entropy maximisation, the dynamical Hamiltonian constitutes a minimal, essentially assumptionfree model (35, 87, 88).

We show that the dynamics of the inferred architecture are reproduced, including fluctuations and their correlations, by such dynamical Hamiltonians. This establishes that the architecture maps any given sensory input onto a characteristic trajectory that maximises path entropy. In other words, the non-equilibrium dynamics represent sensory inputs as probabilistic patterns of activity change (joint toggling of neural assemblies). The representation is fundamentally robust because it is governed by a variational principle of statistical physics.

Trajectory increments over a short interval Δ*t* are generated by random variables *l*^+^ and *l*^*−*^ that count the number of columns within each population that toggle on or off, respectively. In an architecture with four population activities *e, e*′, *r, r*′, eight independent variables 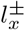 are required (two for each population): 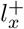 counts inactive columns that switch on and 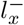 counts active columns that switch off, for *x ∈ {e, e*′, *r, r*′*}*. Each activity *x* is restricted to the integer range *{*0, 1, …, *N*_*x*_*}*. The resulting increments are

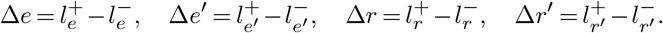

The dynamical Hamiltonian specifies how the probabilities of all possible increments depend on the current population activities:

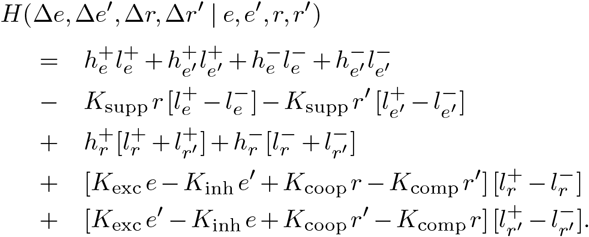

Here *e, e*′*∈ {*0, 1, …, *N*_*e*_*}*and *r, r*′ *∈* {0, 1, …, *N*_*r*_*}* are the instantaneous activities, and the eleven Lagrange multipliers are 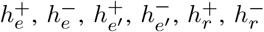, *K*_coop_, *K*_comp_, *K*_exc_, *K*_inh_ and *K*_supp_.

The multipliers 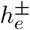 and 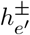 set the basal toggling rates of the lower-level populations and incorporate the effect of the sensory inputs *I* and *I*′. For symmetric input (*I* = *I*′) only nine multipliers need be inferred 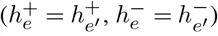. The multipliers 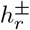 likewise set the basal toggling rates of the upper-level populations. The remaining multipliers encode the interactions: feed-forward excitation and inhibition from lowerto upper-level populations (*K*_exc_, *K*_inh_), cooperative and competitive interactions among upper-level populations (*K*_coop_, *K*_comp_), and feedback suppression from upperto lower-level populations (*K*_supp_). The model retains only terms linear in the dynamical variables.

The probability of an incremental step is then

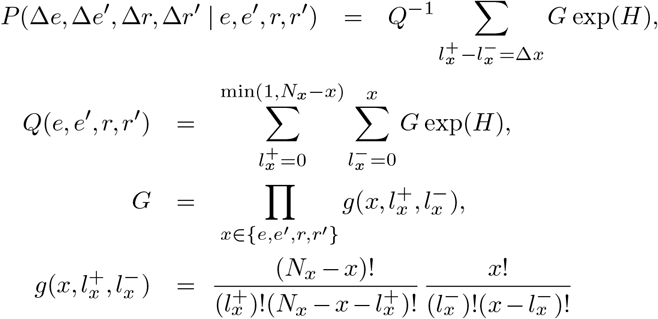

where *Q* is the dynamical partition function and *G* is the degeneracy factor that accounts for the multiplicity of microscopic toggles consistent with a given macroscopic increment.

This construction is formally analogous to, yet fundamentally different from, the canonical ensemble of equilibrium statistical mechanics. It describes non-equilibrium steady states and maximises path entropy over an ensemble of trajectories rather than state entropy over an ensemble of configurations. Consequently the Hamiltonian is not an energy, and the transition probabilities are in general asymmetric:

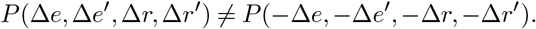

This broken detailed balance is the hallmark of nonequilibrium dynamics. The same formalism has been applied extensively to non-thermal systems, notably to genetic networks in which proteins are produced and degraded (birth– death processes) under feedback regulation (35, 87–92).

We inferred Hamiltonian functions from trajectories generated under two symmetric sensory-input conditions, *I* = *I*′ = 0.25 and *I* = *I*′ = 0.75 (**Fig. 3a**). All trajectories were long enough to sample state space exhaustively and to realise a non-equilibrium steady state, as confirmed by high Hess overlap between half-trajectories (see **Suppl. Fig. 9** and Methods).

**Fig. 3.**
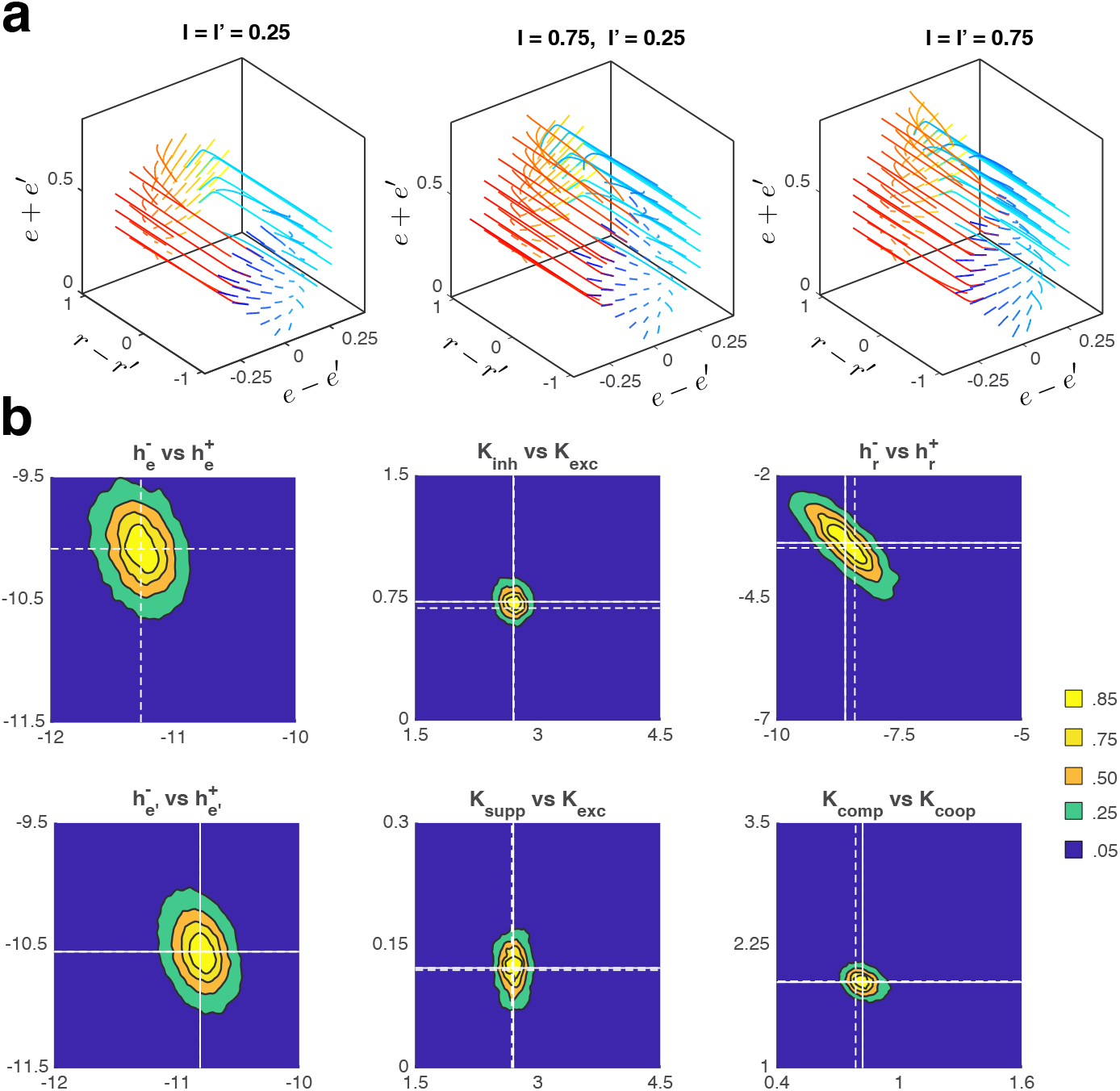
Trajectory flow and fitted Hamiltonian for different input conditions. **(a)** Average trajectory flow for weak symmetric input *I* = *I*′= 0.25 (left), asymmetric input *I* = 0.25, *I*′ = 0.75 (middle), and strong symmetric input *I* = *I*′ = 0.75 (right). Stronger input increases the range of *e* + *e*′ values and shortens (accelerates) trajectory segments during dominance phases. **(b)** Posterior probability distributions (coloured contours) of Lagrange multipliers for symmetric input at *I* = *I*′ = 0.25 and *I* = *I*′ = 0.75. Colour represents the cumulative probability outside each contour. In each panel the cumulative probability is projected onto the plane spanned by two particular multipliers (indicated in the panel title). Multipliers 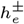 and 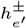 change with input level (0.25 or 0.75), while the remaining multipliers are largely unaffected. White crosshairs indicate the mean values for weak input (*I* = *I* ′= 0.25, dashed) and strong input (*I* = *I*′ = 0.75, solid).

The posterior distributions of the inferred Lagrange multipliers closely resemble nine-dimensional multivariate Gaussians (**Fig. 3b**). These posteriors faithfully reflect the statistics of the non-equilibrium trajectories evoked by each input. They were obtained by standard Markov-chain Monte Carlo sampling for each condition (see Methods). The nine multipliers are largely independent; only a weak trade-off appears between 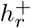 and 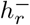, the sum 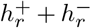 being more tightly constrained than the difference. Input strength affects solely the lower-level multipliers 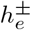 and 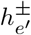; the remaining multipliers are essentially unchanged. This demonstrates that the effects of the four *h*_*e*_ multipliers are largely independent of the interaction terms.

Because the dynamical Hamiltonian is a minimal model, it can only approximate the master-equation dynamics. Nevertheless, the correspondence between Hamiltonian-generated and master-equation trajectories is quantitatively excellent. We verified this with several independent metrics—Hess overlap, differential trajectory flow, differential trajectory dispersion, and the characteristic statistics of dominance reversals (see **Suppl. Figs. 9–11** and Methods). The close agreement confirms that the mesoscale dynamics conform to the variational principle of maximum caliber.

To demonstrate generality we also inferred Lagrange multipliers from the dissimilar trajectories produced by eight modified models in which the coupling strengths were scaled by factors 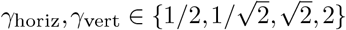 (see Section 2.4). The resulting posteriors are shown in **Suppl. Figs. 12–13**. Scaling the couplings changes the corresponding multipliers approximately linearly while leaving the others essentially unaltered. This establishes the mutual independence of *K*_coop_, *K*_comp_, *K*_supp_, *K*_exc_ and *K*_inh_. The close match between original and Hamiltonian trajectory flows is illustrated in **Suppl. Fig. 9b**.

### Role of small-number fluctuations

It is not obvious why a perceptual system would rely on birth–death processes that generate internal noise in the form of stochastic on–off toggling. At first glance one might expect such internal noise to be detrimental to function, since it would compound any external noise present in the sensory inputs. The functional cost would depend on the relative strength of internal and external noise, which is difficult to know a priori. On further reflection, however, internal noise could also benefit function by making the dynamics more flexible and more sensitive to external inputs. The underlying intuition is that intrinsic noise may enable a form of stochastic resonance, thereby improving the system’s ability to detect or respond to weak external signals that would otherwise remain subthreshold. The functional importance of noise has been established in many biological systems, including translational bursting, gene switching, excitable membranes, vesicular transport and synaptic transmission (93–99).

The reversal statistics of human perception constrain the underlying architecture to small, discrete populations. Approximately two dozen toggling components per population (*N*_*e*_ *≈* 25) quantitatively reproduce both the shape of dominanceduration distributions and the sequential correlations that increase with input strength (see **Fig. 2d**). Slightly larger populations (e.g. *N*_*e*_ *≈* 40) reproduce these features significantly less well (49). Below we show that smaller populations confer a surprising functional benefit: they largely eliminate trajectory “inertia”. At the same time they impose a functional cost by reducing the rate of information uptake from sensory inputs.

To clarify the role of small-number fluctuations we examined the inferred architecture across a range of population sizes *N*_*e*_ and *N*_*r*_ (**Fig. 4**). Fractional activities vary binomially, so that the standard deviation of spontaneous fluctuations scales as 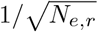. We investigated *N*_*e*_ *∈* {25, 50, 100, 250, 500, 1000, *∞}*and *N*_*r*_ *∈* {10, *∞*}. For this comparison we used the three-alternative architecture, in which every dominant state has two possible successors and trajectories may therefore turn in either direction. (In the two-alternative case dominance simply alternates between the two states.)

**Fig. 4.**
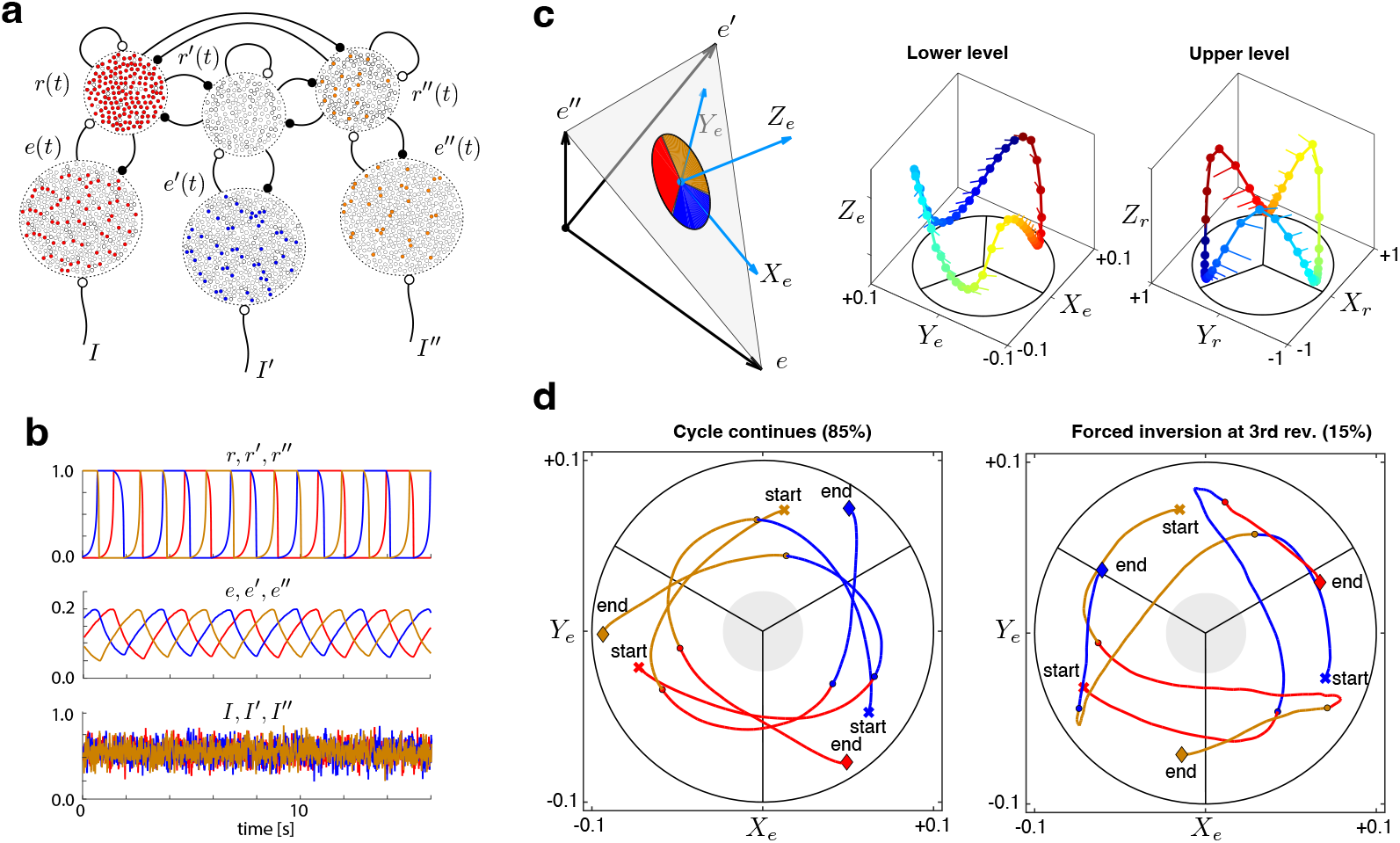
Deterministic dynamics in the diffusion limit. **(a)** Three-alternative architecture with large toggling populations (schematic). Dynamics were examined for *N*_*e*_ = 25, 50, 100, 250, 500, 1000 and *∞*. **(b)** In the diffusion limit *N*_*e*_, *N*_*r*_ *→∞*, trajectories become deterministic; cycle direction is set by initial conditions. This trajectory inertia persists even in the presence of external noise (here *I, I*′, *I*″ = 0.5 *±* 0.1). **(c)** The six-dimensional state space (lower-level activities *e, e*′, *e*″ and upper-level activities *r, r*′, *r*″) can be visualised in three-dimensional subspaces using simplex coordinates (*X*_*e*_, *Y*_*e*_, *Z*_*e*_) and (*X*_*r*_, *Y*_*r*_, *Z*_*r*_) (left). Coloured sectors indicate the phase angles that correspond to different dominance states. Phase-averaged trajectories of lower-level activity (middle) and upper-level activity (right) are shown, with colours representing phase angle. Note the phase shift between lower and upper levels and its relation to the sector boundaries in the *X*–*Y* plane. Phase flow over 0.1 s (whiskers) indicates cycle direction; both directions are equally possible. **(d)** Strong external noise that oversaturates the normal dynamic range [0, 1] can occasionally invert cycle direction (here *I, I*′, *I*″ = 0.5 *±* 0.5). Average lower-level trajectories over three dominance periods, starting from an initial reversal (crosses) and ending after three further reversals (diamonds). Colours indicate the concurrent upper-level dominance state. Despite strong forcing, 85% of third reversals continue the established cycle direction (left) while 15% invert it (right). All trajectories remain well outside the ambiguous zone (grey disk).

#### Large populations loose responsiveness

In the diffusion limit *N*_*e*_, *N*_*r*_*→∞* the master equations become deterministic and sensory input remains the only source of noise (see Methods). Trajectories are then predictable and dominant states succeed one another in a fixed cyclic order (e.g. *r →r*′ *→r*″ *→r*). Because the equations are symmetric, the opposite cycle direction is equally admissible; symmetry is broken only by initial conditions, which select the prevailing limit cycle. Certain initial conditions produce limit cycles that alternate between only two of the three states. Trajectory inertia is strong: once established, a limit cycle tends to persist even in the presence of external sensory noise.

The origin of this inertia is hysteresis. When upper-level dominance switches, for example from *r* to *r*′, feedback inhibition suppresses the corresponding lower-level activity *e*′ while *e* recovers. The third lower-level activity *e*″, however, has already begun to recover during the preceding period and therefore enjoys a head-start. Consequently the next reversal is biased from *r*′ toward *r*″ (**Fig. 4b**).

The state space of the three-alternative system is sixdimensional. Limit-cycle trajectories can be visualised in the three-dimensional subspaces spanned by the lower-level or upper-level activities (**Fig. 4c**). In each subspace the three population activities rise and fall in alternation while total activity remains approximately constant. A convenient coordinate system is the simplex plane defined by the points (1, 0, 0), (0, 1, 0) and (0, 0, 1). In simplex coordinates (*X, Y, Z*), position in the *X*–*Y* plane encodes relative activity and position along the *Z*-axis encodes average activity. Trajectories circulate about the *Z*-axis; each dominance state occupies a characteristic range of phase angles (coloured sectors). The *Z*-axis and its immediate neighbourhood constitute a zone of ambiguity in which differential activity vanishes.

A representative limit-cycle trajectory is shown in **Fig. 4c**. Relative lower-level activity changes continuously while average activity is modulated periodically (long-term mean ~ 0.22). The same holds for upper-level activity (long-term mean ~ 0.45). Lower-level phase angles lead upper-level phase angles by approximately 3*π/*4. Reversals of dominance are preceded by a peak in the lower-level trajectory and a trough in the upper-level trajectory. Cycle direction is set by initial conditions; in the illustrated example the trajectory progresses clockwise (whiskers).

When sensory noise is sufficiently strong the cycle direction can invert (see also **Suppl. Fig. 14**). **Fig. 4d** shows average trajectories spanning three successive dominance periods. Sensory inputs were independent Ornstein–Uhlenbeck processes with mean *µ*_*I*_ = 0.5, standard deviation *σ*_*I*_ = 0.5 and autocorrelation time *τ*_*I*_ = 0.1 s. Most trajectories (85%) continued in the established direction, while a minority (15%) inverted at the third reversal. In both cases the trajectories remained well outside the zone of ambiguity (grey disk). With weaker noise the inversion probabilities fell to 2.4 *±* 0.4% (*σ*_*I*_ = 0.25) and 0.0 *±* 0.2% (*σ*_*I*_ = 0.1).

#### Small populations preserve responsiveness

Dynamics with small populations are qualitatively different. Toggling noise alone causes the cycle direction to change frequently and spontaneously, even under constant sensory input (**Fig. 5a**). Because trajectories proceed in both clockwise and counterclockwise directions with comparable frequency, no single cycle direction prevails. Phase-averaged trajectories of lowerand upper-level activity are shown in **Fig. 5b**. Residual flows point radially toward the origin because circumferential components in opposite directions cancel.

**Fig. 5.**
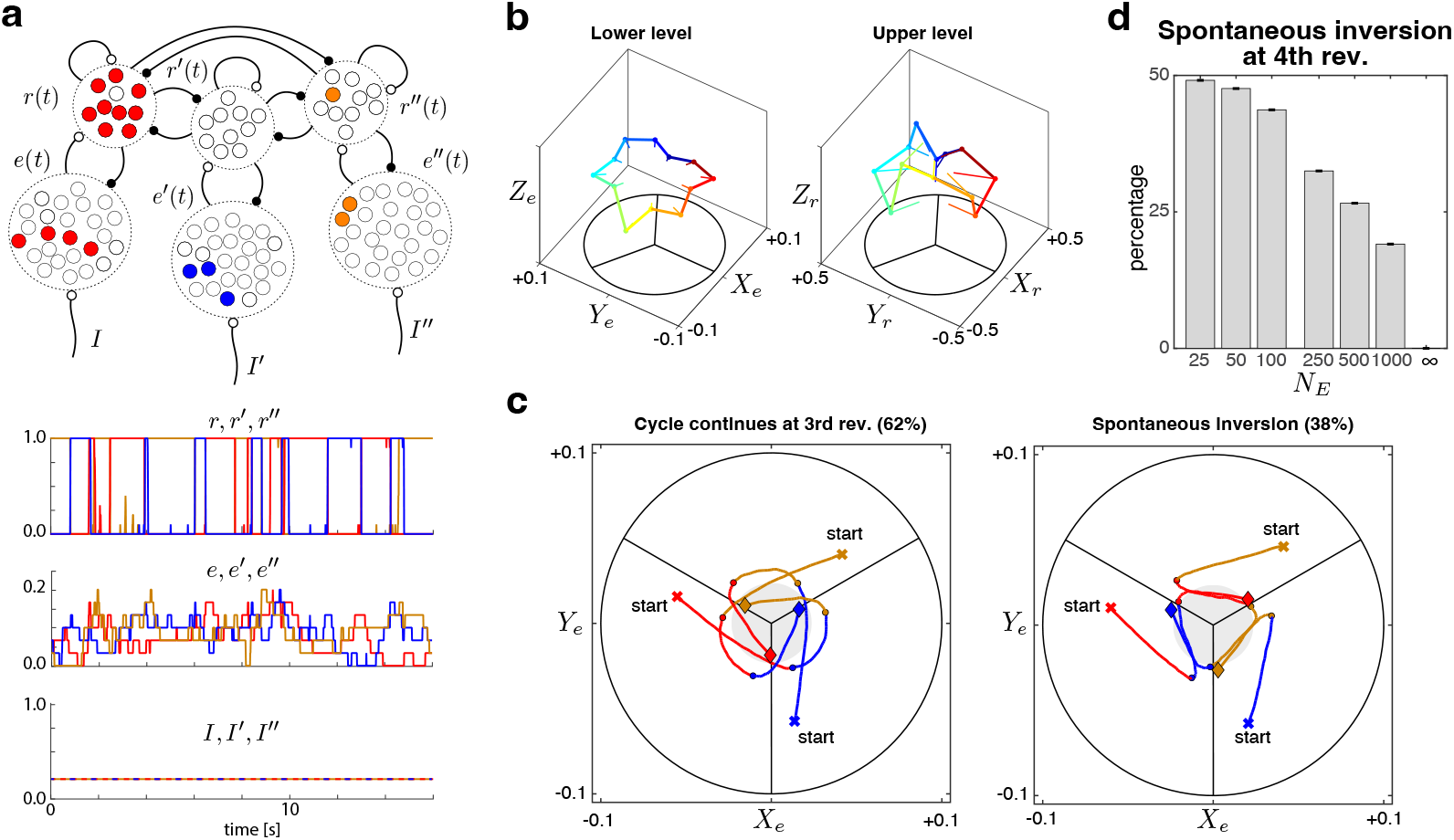
Stochastic dynamics of the discrete system. **(a)** Three-alternative architecture with *N*_*e*_ = 25 and *N*_*r*_ = 10 (top). Toggling noise frequently inverts cycle direction (bottom) and nearly eliminates trajectory inertia, even under constant input (*I* = *I*′ = *I*″ = 0.25). **(b)** Phase-averaged trajectories of lower-level activity (*X*_*e*_, *Y*_*e*_, *Z*_*e*_) (left) and upper-level activity (*X*_*r*_, *Y*_*r*_, *Z*_*r*_) (right) for twelve phase sectors (colours). Residual flow (whiskers) is radial toward the origin because circumferential flows in opposite directions cancel. **(c)** Representative average lower-level trajectories over three dominance periods, starting from an initial reversal (crosses) and ending at a third subsequent reversal (diamonds). Colours indicate upper-level dominance state. Only 62% of third reversals continue the established cycle direction (left); 38% invert it (right). Although trajectories begin farther from the origin, they subsequently approach the ambiguous zone (grey disk). **(d)** Probability of spontaneous trajectory inversion at the fourth reversal for different *N*_*e*_, always with *N*_*r*_ = 10. Only for *N*_*e*_ = 25 is the cycle direction is nearly random. With increasing population size the dynamics become progressively more deterministic; in the infinite-population (diffusion) limit, cycle direction persists indefinitely.

Representative average trajectories spanning three dominance periods are illustrated in **Fig. 5c**. Although the displayed trajectories are counterclockwise, clockwise trajectories occur equally often. Trajectories begin at some distance from the origin (where activity differences are large) but subsequently approach the ambiguous zone (where differences are small). This central tendency contrasts with the behaviour observed in the diffusion limit. After two successive periods in one direction the next reversal continues that direction with probability 62 *±* 0.1% and inverts it with probability 38 *±* 0.1%; by the fourth reversal the inversion probability rises to 49*±*0.1%. Thus spontaneous toggling of only *N*_*e*_ = 25 bistable columns per population largely eliminates trajectory inertia—even in the complete absence of external noise.

To quantify the dependence on population size we measured the probability of cycle-direction inversion at the fourth reversal for *N*_*e*_ *∈* {25, 50, 100, 250, 500, 1000} (with *N*_*r*_ = 10 fixed; **Fig. 5d**). Significant persistence of the established direction is observed for all sizes except the smallest (*N*_*e*_ = 25). We conclude that bistable toggling of sufficiently small populations confers a crucial functional benefit on circuit architectures that rely on strong intrinsic interactions for non-linear amplification. Without toggling noise the dynamics of such architectures become dominated by their recent history and can become trapped in limit cycles, losing responsiveness to external input. Bistable toggling counteracts this undesirable side-effect of strong interactions and ensures responsiveness to input.

### Interactions maximize trajectory entropy

Perceptual evidence constrains not only the size of the toggling populations but also the strength of the interactions among them. In particular, it constrains the reciprocal (cooperative and competitive) couplings among upper-level populations as well as the non-reciprocal (stabilising and destabilising) couplings between lower- and upper-level populations. We asked whether the non-equilibrium steady state (NESS) generated by these interactions is distinguished in any principled way—for example, by its distance from equilibrium. To address this question we compared a family of partially dissimilar NESS dynamics and quantified several measures of non-equilibrium, including entropy production, irreversible flow and trajectory caliber.

Dissimilar NESS dynamics were generated by scaling the coupling strengths with multiplicative factors *γ*_horiz_ and *γ*_vert_ drawn from the set 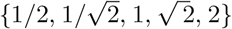. To limit the combinatorial possibilities we scaled couplings in two groups: “horizontal” interactions within the upper level (*w*_coop_ and *w*_comp_) and “vertical” interactions between upper and lower levels (*w*_exc_, *w*_inh_ and *w*_supp_). Horizontal interactions govern dynamic stability via the potential landscape of the upper level, whereas vertical interactions control dynamic sensitivity by amplifying external input differentials. Stability and sensitivity are conflicting objectives that are balanced by multistable perception (100).

Scaling the horizontal and vertical interactions produced nine distinct dynamical regimes (**Fig. 6a**). Each NESS is identified by a pair (*γ*_horiz_, *γ*_vert_) and is illustrated by short average trajectories in the three-dimensional subspace (*e − e*′, *e* + *e*′, *r − r*′). All regimes alternate between dominance phases (stable *r, r*′ with slowly varying *e, e*′) and reversal phases (rapidly varying *r, r*′). The time spent in each phase, however, differs markedly: stronger horizontal and weaker vertical interactions enhance stability and lengthen dominance phases, whereas the opposite combination enhances sensitivity and produces more frequent reversals.

**Fig. 6.**
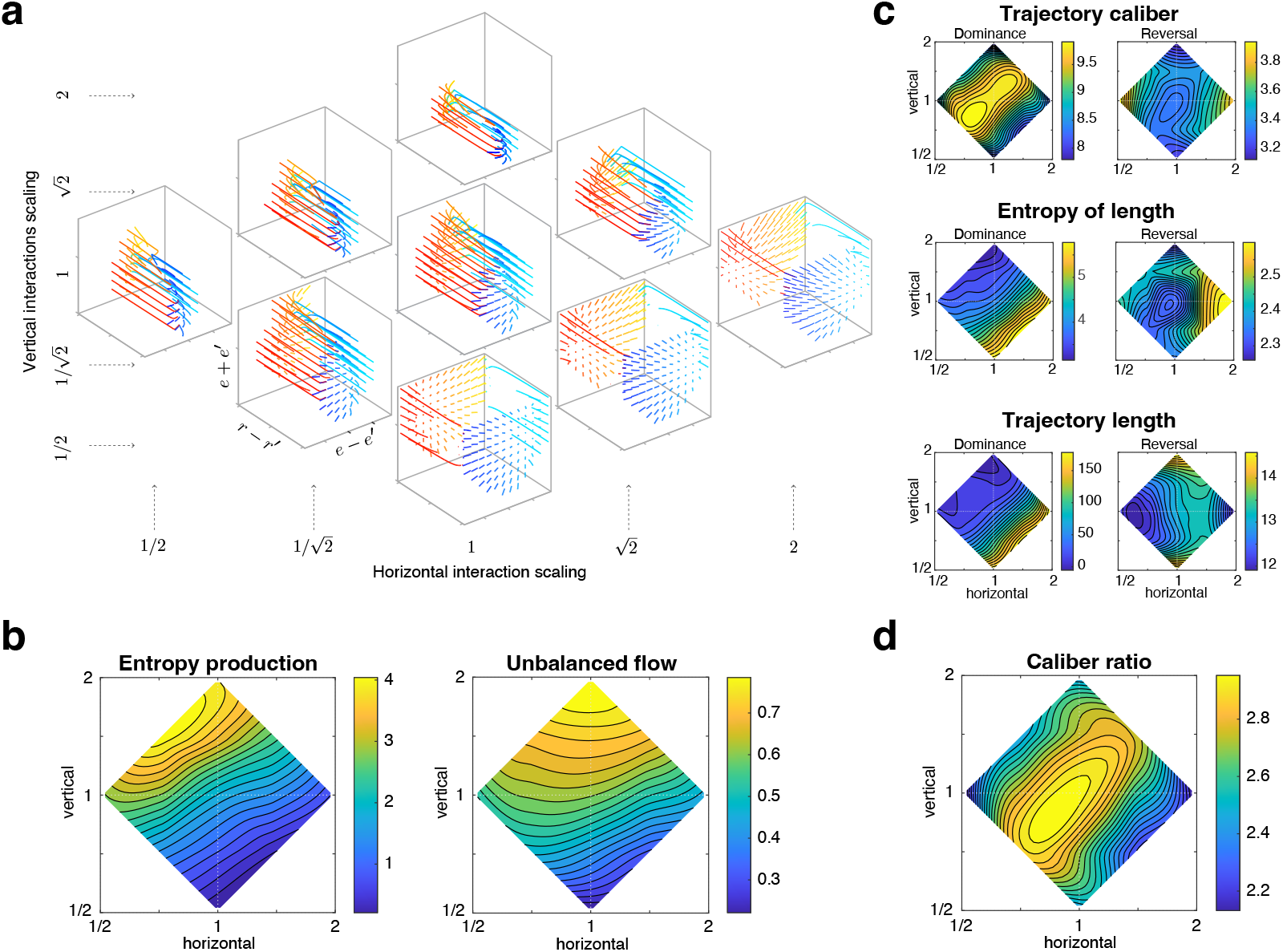
Comparison of dissimilar NESS dynamics. **(a)** Nine NESS dynamics obtained by multiplicatively scaling horizontal and vertical interactions with factors *γ*_horiz_ and *γ*_vert_. The perceptual NESS lies at the centre (*γ*_horiz_ = *γ*_vert_ = 1). Each dynamic is illustrated by the average trajectory flow in the subspace (*e − e*′, *r − r*′, *e* + *e*′). During dominance phases *r− r*′ remains nearly constant (*±*1) while *e* and *e*′ change slowly; during reversal phases *r − r*′ changes rapidly. **(b)** Distance from equilibrium measured by entropy production *S* (left) and by the fraction of unbalanced flow NF*/*TF (right). Contour plots were obtained by interpolation among the nine simulated points. **(c)** Trajectory caliber (top), entropy of trajectory length (middle) and average trajectory length (bottom), shown separately for dominance phases (left) and reversal phases (right). **(d)** Ratio of trajectory caliber in dominance versus reversal phases. The perceptual NESS at the centre combines maximal caliber during dominance with minimal caliber during reversals.

To quantify these dynamics we compared several measures of distance from equilibrium (see Methods). A standard measure is entropy production 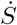. For any pair of states *i, j* with forward and reverse flows *f*_*ij*_ and *f*_*ji*_, the contribution to entropy production is 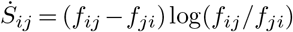. This quantity underestimates the true entropy production of a NESS because only pairs with non-zero flows can be included (to avoid singularities).

An alternative measure is the total net flow,

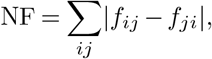

normalised by the total flow

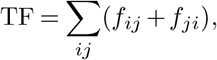

both summed over all pairs of states. The ratio NF*/*TF counts all asymmetries, including pairs in which one of the flows vanishes.

Results for entropy production and the net-flow ratio are shown as contour plots in **Fig. 6b**; the good/bad-actor ratio yielded essentially identical patterns and is omitted. Contours were obtained by interpolating the nine discrete measurements in the (*γ*_horiz_, *γ*_vert_) plane. The centre location corresponds to the NESS of the standard model inferred from perception; the surrounding locations represent dynamics generated with scaled interaction strengths. Both measures give qualitatively similar results: distance from equilibrium increases mostly with the strength of vertical interactions, while horizontal interaction strength has a smaller effect. Thus distance from equilibrium does not single out the inferred perceptual NESS.

We next analysed the dominance and reversal phases of the NESS cycle separately, because the two phases are affected differently by *γ*_horiz_ and *γ*_vert_. A principled measure of trajectory variability is path entropy (caliber), which enumerates all observed trajectory variants (**Fig. 6c**, top row). During dominance phases caliber is maximal when vertical and horizontal interactions are scaled equally (main diagonal of the left panel). During reversal phases caliber is minimal when horizontal interactions are of intermediate strength (vertical midline of the right panel).

These patterns are specific to caliber. Neither the entropy of trajectory length (middle row) nor the average trajectory length (bottom row) exhibits comparable trends. During dominance phases the entropy of length simply tracks average length; during reversal phases it shows only a shallow minimum near the perceptual NESS, indicating greater trajectory consistency.

Consequently the perceptual NESS is distinguished by maximally variable trajectories during dominance phases and maximally stereotypical trajectories during reversal phases. In other words, caliber is maximised while the system is susceptible to sensory input (i.e., while sensory evidence can be evaluated) and minimised while dominance is being reversed (i.e., while a decision is being taken). This organisation is summarised in **Fig. 6d**, which shows that the ratio of dominance-phase to reversal-phase caliber peaks near the perceptual NESS at the centre of the plane.

Greater variability (higher caliber) arises because trajectories linger near the ambiguous zone (see **Fig. 5c**), where lowerlevel activities are comparable and the number of possible continuations is larger. This property is likely to be advantageous for decision-making in volatile environments: changing external input can more readily steer a trajectory that lies close to the ambiguous zone than one that lies far from it. We conclude that the perceptual NESS is functionally organised so as to render trajectories more sensitive to external sensory input, thereby facilitating flexible decision-making under volatile sensory conditions.

### Predicted spiking activity in sensory cortex

As we have seen, the dynamics of the mesoscale model are non-equilibrium steady states (NESS) characterised by unbalanced trajectory flows. Our analyses so far have taken advantage of the fact that model simulations provide complete visibility into the time evolution of every state variable (every column of every population). Experimental recordings, by contrast, afford at best partial visibility—for example, simultaneous recordings from only a few cortical columns.

Here we use the NESS hypothesis to derive predictions that can be tested in sensory cortex with current neurophysiological methods. We subjected the three-alternative architecture to time-modulated input and simulated the evoked response in terms of the toggling activity of a small number of columns: two columns each from the upper-level populations *r, r*′, *r*″ and the lower-level populations *e, e*′, *e*″ (**Fig. 7a**). From these 12 columns we generated Poisson spike trains for 10 neurons per column (120 neurons in total). All subsequent analyses were based on 100 trials of 6 s each, yielding 600 s of total recording time—a realistic volume of data for contemporary cortical recordings.

**Fig. 7.**
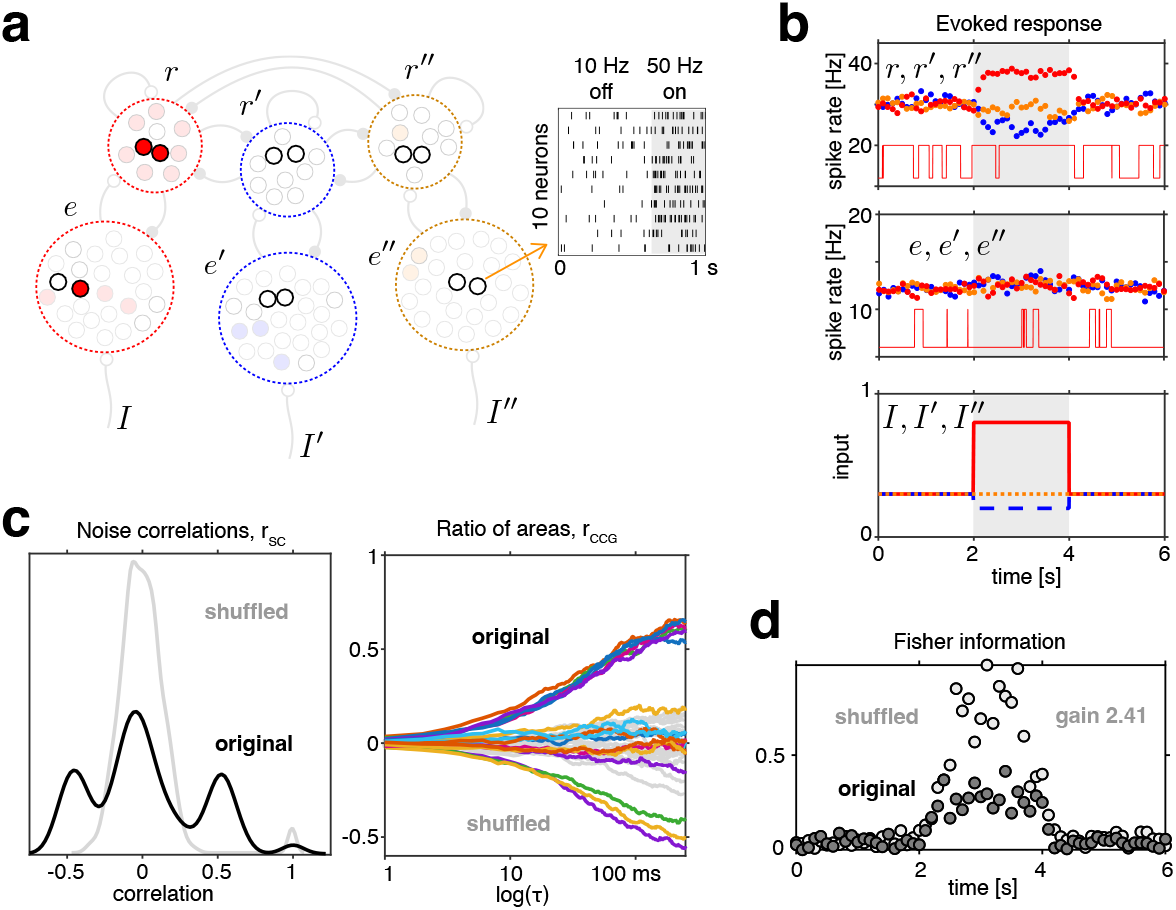
Predicted neuronal responses and correlations. **(a)** On–off toggling of selected columns (two columns per population, 12 columns in total) was used to generate Poisson spikes for 10 neurons per column (120 neurons altogether). Representative spiking from one column is shown in the inset. Spike rates were 10 Hz in the off state and 50 Hz in the on state. **(b)** Average firing rates evoked by differential modulation of *I, I*′ and *I*″ (bottom). Rates (averaged over 20 neurons and 100 trials) are shown in 100 ms bins for lower-level populations *e, e*′, *e*″ (middle) and upper-level populations *r, r*′, *r*″ (top). Representative toggling of one *e*- and one *r*-column in a single trial is indicated by square waves. **(c)** Positive and negative correlations among pairs of upper-level neurons. Left: distribution of noise correlations *r*_SC_ (trial-by-trial spike-count fluctuations). Positive peaks correspond to pairs within the same column or population; the negative peak reflects mutual inhibition between populations. Right: ratio of areas *r*_CCG_ under cross-versus auto-correlograms as a function of lag *τ* for a random subset of pairs. Both positive and negative correlations are evident for *τ <* 200 ms. Random time-shifting abolishes all structure (grey). **(d)** Encoding of alternative input modulations by the 60 upper-level neurons. Fisher information about alternative inputs is shown in 100 ms bins. Random time-shifting removes information redundancy and more than doubles encoding capacity (grey symbols).

#### Spiking simulations

Spiking activity was generated with instantaneous Poisson rates of 10 Hz or 50 Hz according to whether a column was in the off or on state, respectively. Mean toggling rates were 2.9 Hz for lower-level columns and 3.5 Hz for upper-level columns. Dominance episodes of upper-level populations lasted on average 0.48 *±* 0.35 s.

Trial-averaged responses to time-modulated input are shown in **Fig. 7b**. Baseline input was *I* = *I*′ = *I*″ = 0.3. Between 2 s and 4 s one input was raised to *I* = 0.6 (red), another was lowered to *I*′ = 0.2 (blue), and the third remained unchanged at *I*″ = 0.3 (orange). For each population the average firing rate of 20 neurons (drawn from two columns) is plotted in 100 ms bins. Upper-level columns *r, r*′, *r*″ showed clear rate modulation, whereas lower-level columns *e, e*′, *e*″ were affected only marginally. The reason is that lower-level on states are typically brief and intermittent, as illustrated by the representative toggling time-course of a single column in one trial (red square wave). Average rates reflect 200 such time-courses (two columns *×* 100 trials).

#### Predicted correlation structure

Because neuronal spiking is modulated by column toggling, the resulting variability is shared among neurons belonging to the same column and, to a lesser extent, among neurons in different columns of the same population or even of different populations. This shared variability produces pairwise correlations, illustrated in **Fig. 7c** for the 60 upper-level neurons (1770 pairs). Noise correlations *r*_SC_ were computed from trial-by-trial fluctuations in spike counts over a 500 ms window (2.0–2.5 s). The distribution of *r*_SC_ (left panel) exhibits several distinct peaks: a near-unity peak for neuron pairs within the same column, a secondary positive peak (*ρ ≈* 0.5) for pairs in different columns of the same population, and a negative peak (*ρ ≈ −*0.4) reflecting mutual inhibition between populations. Correlations vanish when spike trains are randomly timeshifted (pale distribution).

The temporal structure of shared variability is revealed by cross-correlograms. A convenient summary is the ratio of areas *r*_CCG_ under the cross- and auto-correlograms (101), shown for 17 randomly selected pairs (**Fig. 7c**, right). This measure asymptotes to *r*_SC_ at large lags. Both positive and negative correlations are evident on a characteristic timescale of approximately 100 ms. Random time-shifting of the spike trains abolishes the structure (pale traces).

The spiking activity of the 60 upper-level neurons encodes information about the input modulation. Fisher information about alternative inputs was computed from the trialaveraged responses and their covariance in 100 ms bins (see Methods). Results are shown in **Fig. 7d** (dark symbols). Shared variability introduces redundancy and therefore reduces Fisher information; randomly time-shifting the spike trains removes this redundancy and more than doubles the encoded information (pale symbols).

#### Predicted arrow-of-time irreversibility

In the circuit architecture of the NESS hypothesis some populations are coupled non-reciprocally while others are coupled reciprocally (**Fig. 8a**). Non-reciprocal coupling predicts unbalanced (non-equilibrium) trajectory flows; reciprocal coupling predicts balanced (equilibrium) flows. We tested these predictions by computing arrow-of-time irreversibility, a physics-inspired measure of non-equilibrium flow (102). One hundred trials were concatenated to yield 600 s of quasi-continuous spiking activity (including both modulated and unmodulated epochs). For each of the six populations we obtained the average firing rate of 20 neurons (two columns) in 10 ms bins.

**Fig. 8.**
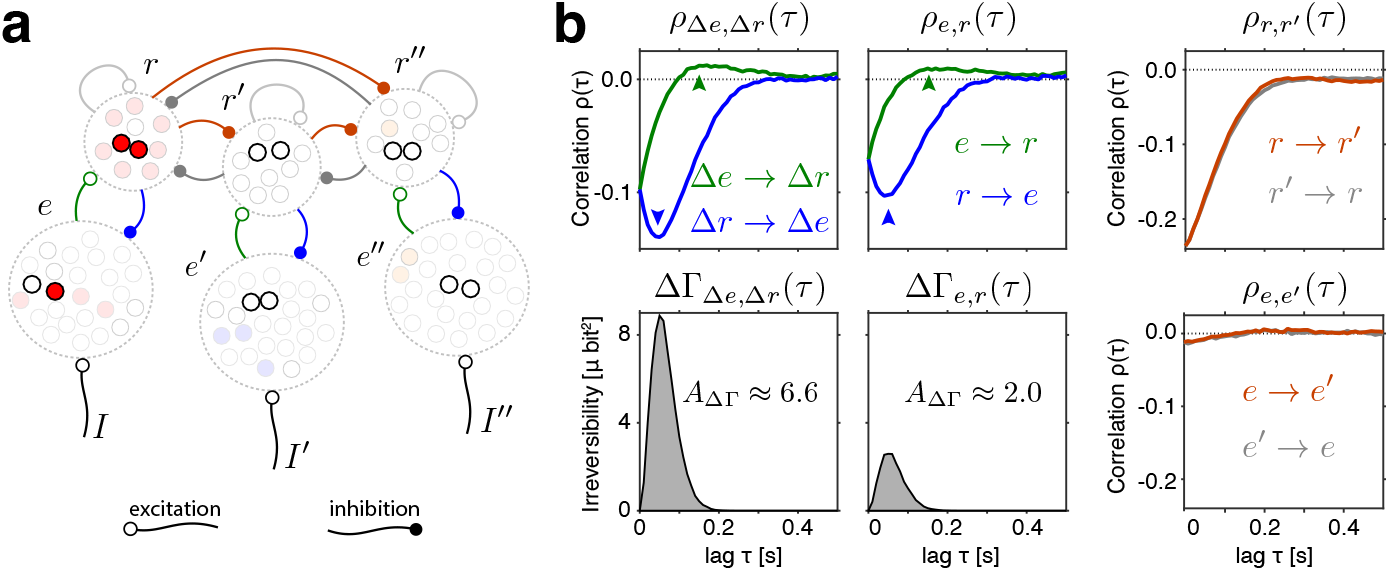
Delay-dependent correlations and arrow-of-time irreversibility. **(a)** Circuit architecture with non-reciprocal and reciprocal couplings. Couplings between lower-level populations (*e, e*′, *e*″) and upper-level populations (*r, r*′, *r*″) are non-reciprocal: feed-forward excitation (green) and feedback inhibition (blue). Couplings among upper-level populations are reciprocal (mutual inhibition; brown/grey). **(b)** Lagged correlations *ρ*(*τ*) between pairs of populations, computed from the firing rates of 20 neurons (two columns) per population in 10 ms bins, concatenated over 100 trials of 6 s (600 s total). Between levels the lagged correlations are asymmetric (top left and middle), producing significant arrow-of-time irreversibility ΔΓ(*τ*) (bottom left and middle). Within levels the correlations are symmetric (right) and irreversibility is negligible (not shown). The asymmetry is especially clear for differential activities Δ*e* = *e −* (*e*′ + *e*″)*/*2 and Δ*r* = *r −* (*r*′ + *r*″)*/*2. The excitatory influence *e →r* is weaker and slower (small peak near 150 ms, green arrows) than the inhibitory influence *r →e* (larger peak near 50 ms, blue arrows).

Arrow-of-time irreversibility ΔΓ(*τ*) is defined from a comparison of lagged pairwise correlations in the forward and reverse directions. Let *ρ*_*x → y*_(*τ*) and *ρ*_*y → x*_(*τ*) denote the lagged correlations of the time-varying activities *x*(*t*) and *y*(*t*) for lag *τ ≥* 0. After conversion to signed mutual information 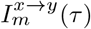 and 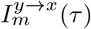, the lag-dependent irreversibility is the squared difference

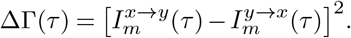

The integrated area *A*_ΔΓ_ = ∫ΔΓ(*τ*) *dτ* quantifies total irreversibility and causal influence.

Predictions are shown in **Fig. 8b**. Non-reciprocal coupling between lowerand upper-level populations (feed-forward excitation *e →r* and feedback inhibition *r →e*) produces lagged correlations that are asymmetric. The correlation *e →r* exhibits a small positive peak near 150 ms, reflecting the weakly excitatory influence of *e*-column toggling on *r*column toggling. Conversely, the correlation *r →e* exhibits a larger negative peak near 50 ms, reflecting the stronger inhibitory influence of *r*-columns on *e*-columns. The difference in strength arises because *e*-toggles typically occur singly, whereas *r*-toggles are typically collective. The observed delays reflect the fact that populations are coupled through toggling kinetics: although a source population alters the toggling rates of its targets instantaneously, relaxation to the new steady state requires time. The characteristic relaxation time of a birth–death process is not fixed but depends on both the current state and the current input (69).

The effects of non-reciprocal coupling become even clearer when differential activities Δ*e* = *e −* (*e*′ + *e*″)*/*2 and Δ*r* = *r −* (*r*′ + *r*″)*/*2 are examined (top-left panel of **Fig. 8b**). Lagged asymmetry ΔΓ(*τ*) is substantially larger for these differential signals than for the individual activities *e* and *r* (bottom-left and middle panels). In both cases the asymmetries are highly significant relative to randomly time-shifted controls (not shown).

Reciprocal coupling (mutual inhibition) among upper-level populations *r* and *r*′ produces strong negative correlations that are symmetric; no significant difference exists between *r →r*′ and *r*′*→r* (top-right panel). Lower-level populations *e* and *e*′ are coupled only indirectly through their corresponding upper-level populations and therefore exhibit only weak, symmetric negative correlations (bottom-right panel). Relative to randomly shifted controls, lagged asymmetry ΔΓ(*τ*) does not reach significance (not shown).

These results provide detailed, quantitative predictions of the NESS hypothesis. Column toggling introduces shared variability in neuronal firing that manifests both as noise correlations and as peaked cross-correlograms. Positive correlations among neurons within the same column, together with characteristic correlation times of order 50 ms, would support the hypothesis of column-level toggling. Non-reciprocal coupling between lowerand upper-level columns produces unbalanced flows and measurable arrow-of-time irreversibility. Observation of irreversibility between cortical levels, but not within levels, would constitute evidence for non-equilibrium dynamics.

## Discussion

Brain activity arises from interactions across different spatial scales, including the *microscale* of individual neurons, the *mesoscale* of local assemblies of neurons, and the *macroscale* of brain systems. Computational models of these interactions can be constructed ‘directly’ (bottom-up from known components), ‘inversely’ (top-down from observed activity or behaviour), or ideally both (103, 104). Functional theories—how interactions between scales generate perception and cognition—remain fragmentary, and substantive further progress may require a paradigm shift.

Here we revisited an inverse model (49) with several interesting and unusual features. First, the model’s point of departure is the macroscale dynamics of perception as experienced and reported by human observers. This starting point provides a direct window onto the neural levels that drive perceptual function. Second, constrained by the empirical evidence, the model posits *discretely* stochastic and *non-equilibrium* dynamics of the underlying neural activity—specifically, populations of ‘toggling’ columns. Third, the model can potentially be grounded at the *mesoscale* of the sensory cortex, in that the inferred components (toggling columns and their interactions) echo the known neural activity and architecture at this level.

### Representing sensory input with flows

The mesoscale model responds to constant input with a particular nonequilibrium steady state (NESS) that is specific to this input. The trajectory is stochastic and exhibits large variation between individual episodes of ‘dominance’ and ‘reversal’. Nevertheless, the distribution of incremental trajectory steps is fully determined by the input and maximises *path* entropy in accordance with the variational principle of nonequilibrium (34, 35). We have shown that this step distribution is reproduced by a dynamical Hamiltonian function and its Lagrange multipliers.

While the step distribution itself has a complex structure, it can be visualised—indirectly but accurately—in terms of another distribution of simpler structure, namely the posterior distribution of Lagrange-multiplier values. This posterior distribution, which was obtained with Markov-Chain Monte Carlo and approximates a multivariate Gaussian, serves to illustrate the degree to which a NESS trajectory is representative for sensory input.

A NESS trajectory unfolds in the multivariate state space of joint activities (of lowerand upper-level populations) and differs fundamentally from accepted views of cortical representation. In those views, the neural representation of a stimulus is the stationary pattern of activity obtained by averaging over many trials (a scalar for one ‘response channel’ and a vector for multiple ‘response channels’) (36). Averaging is needed because the responses on individual trials vary substantially and stochastically, making them difficult to interpret (101).

In contrast, we suggest that the confusing variability of trial responses conceals a consistent underlying structure, namely, a particular multivariate NESS trajectory and path probability that could, at least in principle, be specified by a dynamic Hamiltonian function. A central role of multivariate trajectories in neural representation and computation is supported by a growing body of theoretical and modelling work on spiking neural networks (40, 42) and empirical recordings from primate motor cortex (41, 43).

### Indications of functional organization

Several features of the mesoscale model suggest organization for purposes of perceptual function. The most salient are far-from-equilibrium operation, small population size, and the relative strength of interactions between populations. We now discuss these features in turn.

#### Far-from-equilibrium operation

The toggling populations of the mesoscale model constitute birth–death processes (105), with lower-level populations operating mostly near depletion (most columns off). This far-from-equilibrium regime is consistent with perceptual evidence and reproduces the empirical distribution of intervals between perceptual reversals. In addition, this regime is functionally optimal in that it maximises information gain about sensory input. To see this, consider the example of a single population of independently toggling columns. When external input to such a population steps up, population activity rises stochastically, accumulating Fisher information about the size of the step. The rate of information gain is proportional to distance from equilibrium, population size *N*_*e*_, and toggling rate *ν*_*e*_. In other words, information gain is maximal when distance to equilibrium is largest—that is, for the first few ‘on’ toggles in a depleted population.

In the mesoscale model, populations of toggling columns interact with each other. If they were isolated, external input would drive the fractional activity of lower-level populations to an equilibrium between 80% and 100%. Due to inhibitiory feedback from upper-level reversals, the average fractional activity of lower-level populations typically remains below 25%. Thus, by keeping lower-level populations far from equilibrium, network interactions maximise information gain about sensory input.

#### Small population size

A puzzling aspect of the mesoscale model is the small size of toggling populations, which is constrained to approximately two dozen columns by perceptual evidence. The mystery is deepened by the fact that smaller populations encode less sensory information, as just mentioned. However, in the context of the mesoscale model, smaller populations have the surprising benefit of ensuring responsiveness to sensory input, because their activity experiences larger stochastic fluctuations. This raises the fascinating possibility that a few dozen columns may constitute an optimal trade-off between being more informative about and less responsive to sensory input.

We studied the effect of population size with the threealternative model, in which every dominant state has *two* possible successor states. At the so-called ‘diffusion limit’, where population size is infinite and the dynamical development is deterministic, trajectories settle into stable limit cycles, visiting alternative dominance states in a particular sequential order. These limit cycles are surprisingly stable in the presence of fluctuating sensory input. The reason for this ‘inertia’ is that intrinsic interactions are stronger than sensory inputs, so that the dynamics are dominated by hysteresis. Only large sensory fluctuations can disrupt an established cycle of dominance states.

We also investigated the model dynamics with larger population sizes but without any sensory noise. In this case, dominance cycles invert occasionally due to toggling noise. The inversion rate depends on population size, and cycle direction becomes fully randomised only for the smallest populations. Thus, the toggling noise of smaller populations almost eliminates the ‘inertia’ of larger populations. A related finding is that the activities of smaller populations remain in a ‘zone of ambiguity’ (close to each other), so that even comparatively small fluctuations can trigger reversals.

To see why smaller populations are more ‘noisy’, one must compare the rates of ‘drift’ (change of expected activity) and of ‘dispersion’ (growth of expected variance) in a birth– death process (70). Since larger populations decrease dispersion (while leaving drift unchanged), smaller populations are more stochastic.

#### Relative strength of interactions

The mesoscale model includes ‘horizontal’ interactions between upper-level populations as well as ‘vertical’ interactions between lowerand upper-level populations. Broadly speaking, horizontal interactions determine the stability of ‘dominance’ episodes, while vertical interactions control the sensitivity of ‘reversals’ to fluctuations and sensory inputs.

Stability and sensitivity are conflicting objectives of sensory systems. To assess how horizontal and vertical interactions influence the NESS dynamics, we simulated the twoalternative model with nine different combinations of interaction strengths, scaling either horizontal interactions, or vertical interactions, or both. While all of the resulting dynamics alternated between dominance and reversal phases, they were also substantially different. Most obviously, when vertical interactions were stronger relative to horizontal ones, the sensitivity of reversals to lower-level fluctuations was heightened, accelerating the dynamics and shortening dominance episodes. This increased distance from equilibrium, as measured by entropy production or fraction of unbalanced flow. Further differences were evident in the path entropy (caliber) during dominance and reversal phases. Path entropy was obtained by counting how often different trajectories occur. We computed a lower bound by counting the number of increments and decrements in four population activities, disregarding order and timing (see Methods). Path entropy is high when many different trajectories occur comparably often, but low when a few trajectories occur far more often than others. Path entropy also measures the extent to which trajectories are dominated by stochastic fluctuations (dissipation) as opposed to deterministic flow (drift).

Our results show that path entropy during *dominance* phases was *maximal* for the relative strength of vertical and horizontal interactions inferred from perceptual evidence. Both raising and lowering this ratio reduced path entropy. In other words, the ratio inferred from perception maximised the contribution of stochastic fluctuations during dominance phases. We expect that this ratio would also maximise sensitivity to fluctuations originating from sensory input, making it relevant to perceptual function. However, further work will be needed to confirm this expectation.

### Comparison to sensory cortex

There are several intriguing parallels between the mesoscale model and the known architecture and activity of sensory cortex. Before describing these, we briefly summarise the relevant features of sensory cortex. In addition, we discuss the predictions that the mesoscale model makes for spiking activity in sensory cortex.

#### Mesoscale architecture and activity of sensory cortex

In humans, the sensory cortex (visual, auditory, and tactile) covers more than a quarter of the cortical surface, and its micro- and mesoscale structure largely follows a general blueprint (106). This holds also for visual cortex, which covers approximately 25,000 mm^2^ in humans and is organised hierarchically into distinct areas, with ‘higher’ areas typically representing more complex image features and being more invariant to viewing conditions (81, 82). The local connectivity and correlated activity in the cortical sheet delineate mesoscale assemblies with *O*(10^2^–10^3^) nearby neurons (‘cortical columns’) (16, 17, 107, 108). Depending on the definition, there are 100 to 1,000 of these mesoscale columns per mm^2^ of cortex. Within each area, the horizontal connectivity between columns ranges over several millimetres and is typically reciprocal, balancing mutual excitation (between columns with similar inputs) and inhibition (between columns with dissimilar inputs) (109–112). This canonical circuit motif—like excites, unlike inhibits—has a number of important consequences. First, when activity is averaged over trials, responses to image features are normalised ‘divisively’ with respect to other similar features (113–115). Second, the competitive ‘winner-take-all’ dynamic of this circuit creates distinct attractor states of collective activity for nearby columns with similar input (70, 116, 117). Third, fluctuation-driven toggling between these ‘on’ and ‘off’ states can strongly modulate the response on individual trials (18, 118).

The visual cortex is functionally organised in that every column responds to stimulation in a specific part of the visual field (‘receptive field’) (79, 80). The size of receptive fields increases over the hierarchy of cortical areas because ascending projections from distant columns in lower-order areas target nearby columns in higher-order areas (‘convergence’). Between successive levels of the hierarchy, the areal size of receptive fields increases by *O*(10) times. The ‘ascending pathway’ from lowerto higher-order areas originates from superficial layers of cortical columns and is distinct from the ‘descending pathway’ in the opposite direction, which originates from deep layers (81, 119, 120). The targeting of both ascending and descending projections is point-to-point, consistent with directly reciprocal loops between columns (but not between layers) (121, 122). While ascending projections *excite* activity in target columns, the effect of descending projections remains poorly understood (123, 124). A number of computational studies of cortical networks have assumed that descending projections *inhibit* the activity of target columns (119, 125–128).

#### Parallels between model and cortex

The mesoscale model echoes several features of visual cortex. The interaction between upper-level populations corresponds to the competitive winner-take-all dynamic between nearby cortical columns with large and overlapping receptive fields. Lower-level populations with independent units correspond to more distant cortical columns with small and non-overlapping receptive fields. The non-reciprocal projections between lower- and upper-level populations mirror the ascending and descending projections between hierarchical levels of cortex, which connect columns with small and non-overlapping receptive fields to columns with large and overlapping receptive fields (and vice versa). In the model, ascending projections are excitatory, whereas descending projections are inhibitory. Lastly, the stochastic on–off toggling in the model mimics fluctuation-driven transitions between distinct attractor states of local assemblies in cortex.

Because of these parallels, we conjecture that one model circuit with *O*(10) alternatives could be implemented by a small patch of cortex responding to one particular visual location at one level of the hierarchy, together with the connected patches at lower levels of cortex that also respond to this location. The combined cortical area would be approximately 1 mm^2^ with 1,000 cortical columns.

#### Predictions for the microscale of spiking neurons

Although we have implemented mesoscale models with populations of toggling columns, they could also have been realised with assemblies of spiking neurons. Our prior work has shown that stochastic toggling—as well as the input-dependent toggling rates—quantitatively reproduces fluctuation-driven attractor transitions in assemblies of spiking neurons (49, 70). Here we have taken a shortcut and converted column toggling into inhomogeneous Poisson spike trains, assuming differential firing rates during the ‘on’ and ‘off’ states of each column. To generate testable predictions, we simulated an amount of data that could realistically be recorded in a neurophysiological experiment.

#### Shared variability

Our simulations highlight the difference between individual trials and averages over trials. On individual trials, the response is dominated by stochastic toggling between ‘on’ and ‘off’ states, largely obscuring the differences between sensory inputs. In the trial average, the response level is intermediate between ‘on’ and ‘off’ levels and varies with sensory input. This is because the probability of ‘on’ and ‘off’ states is controlled by toggling rates which, in turn, are set by input.

Total spike counts vary from trial to trial even for identical input (because toggling is stochastic). This variability can be correlated positively for neurons in the same population (i.e., with the same stimulus preference), but negatively for neurons in different populations (i.e., with different stimulus preferences). The strength of correlation also depends on sensory input, which determines the probability of ‘on’ states. Specifically, we predict stronger positive correlations within strongly stimulated columns and stronger negative correlations between strongly and weakly stimulated columns. Within or between weakly stimulated columns, the correlations are also weak. These predictions are broadly consistent with ‘noise correlations’ observed in the visual cortex (101).

The correlation structure introduced by column toggling also shapes the cross-correlation of spike times. This can be measured in terms of the cross-correlogram, CCG(*τ*), as a function of lag *τ*, or of the ratio of areas, *r*_CCG_(*τ*), under the cross-correlogram and the auto-correlogram (see Methods and (101)). In our idealised simulations, the ratio *r*_CCG_(*τ*) reaches values as high as *±* 0.5 (for positively or negatively correlated neurons) for delays up to *τ* = 100 ms. In empirical recordings from the visual cortex, spike-time correlations are typically heterogeneous, but can reach similar sizes and durations (18).

#### Arrow-of-time irreversibility

A critical prediction of the mesoscale model is non-equilibrium (unbalanced) flow between cortical columns with non-reciprocal interactions— specifically, columns in lower- and upper-level populations with *similar* stimulus preference. A straightforward measure of non-equilibrium flow is ‘arrow-of-time’ irreversibility, which compares lag-dependent correlations in the forward and reverse directions of time (see Methods). This measure applies to different types of neural activity (5, 75, 102) and reveals both the strength and direction of causal influence.

The mesoscale model predicts a stronger *negative* influence from upper-level to lower-level columns, due to the feedback suppression that is triggered by the largely coordinated toggling of upper-level columns during a reversal. Simultaneously, it predicts a weaker *positive* influence from lower-level to upper-level columns, stemming from the feedforward facilitation by the largely uncoordinated toggling of lower-level columns. Both predictions can in principle be tested neurophysiologically by recording simultaneously from upperand lower-level columns with overlapping receptive fields and similar stimulus preferences. Note that these predictions assume simultaneous recording of multiple neurons from both upperand lower-level columns, so that ‘on’ or ‘off’ states can be distinguished.

### Cortical function and statistical physics

In the broadest sense, brains exploit the lawful structure of natural environments and accumulate past experience to predict the immediate future (129, 130). In visual cortex, this ‘predictive inference’ is thought to be performed hierarchically, with every cortical patch (columns with overlapping receptive fields) at every level of the hierarchy generating predictions about its immediate input (83, 127, 131, 132). Intriguingly, even retinae detached *in vitro* can perform predictive inference for simple stimuli (133, 134).

To explain predictive inference in sensory cortex, some theories have taken inspiration from statistical physics, specifically the variational principle of *equilibrium*—’maximal state entropy’ or ‘minimal free energy’ (47, 135). This attractive hypothesis is summarised below. However, at the scale of the whole brain, neural activity exhibits all the hallmarks of *nonequilibrium* dynamics, such as unbalanced (4–6) or turbulent flows (7, 8, 23). We conjecture that the variational principle of *non-equilibrium* can resolve this apparent conflict and reconcile these positions.

#### Free Energy Principle

Predictive inference could happen spontaneously if cortical activity were governed by a variational principle (135). Sensory cortex is thought to implement a hierarchical ‘generative model’ of the environment— basically a library of alternative ‘causal origins’ for all possible sensory events (47, 129, 136). Given an observed event, such a model could virtually evaluate all possible origins and identify the most likely alternative.

According to current theories, alternative causal origins could be represented by different activity *states*. In every cortical patch at every level of the hierarchy, the immediate input and connectivity would have to shape the potential energy of local activity states so as to reflect the probability of causal origins in the current sensory context (135). In each of these local energy landscapes, activity would perform a noise-driven random walk and gradient descent. The optimal set of causal origins would be identified when free energy has been minimised over the entire hierarchy (the Free Energy Principle) (47, 136).

In practice, a hierarchical system of this staggering complexity—representing all possible causal origins of sensory events—would not equilibrate in finite time, certainly not at the pace at which sensory events are processed. While a major retarding factor would be the interdependence of local landscapes (i.e., every landscape depending on the activity states of others), random walks of activity would compound the problem. For when a new sensory context alters local landscapes, this would register only after activity had explored the new landscapes and converged on new potential wells. Clearly, brains must have evolved ways and means to vastly accelerate convergence.

#### Variational principle of non-equilibrium

The more general variational principle governing both non-equilibrium and equilibrium systems is maximum *path* entropy or ‘Maximum Caliber’ (34, 35). All classical results of both equilibrium and non-equilibrium dynamics can be recovered from Maximum Caliber (e.g., Boltzmann distribution, partition function, Onsager relations).

Importantly, the mathematical formulation of the variational principle of non-equilibrium mirrors that of equilibrium, with path probabilities substituting for state probabilities and dynamical partition functions substituting for static ones. Accordingly, sensory cortex could represent alternative causal origins of sensory inputs in terms of activity *paths*, not states. To make this work, the input and connectivity of cortical patches would have to shape the probability of activity flows—in other words, the ‘topology of conductivity’.

Representing causal origins with paths could dramatically accelerate convergence. When a new sensory context alters the conductivity in a cortical patch, the activity trajectory would change almost instantaneously, so that other cortical patches could register this change after a minimal integration time. In short, the spontaneous flow of cortical activity could optimally infer causal origins simply by following the current paths of maximal conductivity.

## Conclusion

We conclude that spontaneous reversals of perception reflect underlying neural dynamics that are organised functionally so as to be responsive to sensory inputs. The architecture of this mesoscale model echoes both the canonical connectivity of sensory cortex (83, 119, 120) and the hierarchical inference hypothesis mentioned above (47, 135, 136). Competing populations represent alternative causal origins for current sensory inputs. Additional populations accumulate sensory evidence for every causal origin. The interaction between competing and accumulating populations emphasises evidence that is *unexpected* in the sense that it is *not* predicted by the currently dominant alternatives. This emphasis perpetuates the non-equilibrium dynamics, renders all perceptions open to revision, and ensures active updating even without new input. We conjecture that current theories of brain function (the Free Energy Principle) could be reformulated in terms of the variational principle of *non-equilibrium* (Maximum Caliber) and suggest ways of testing these abstract concepts experimentally with the microand mesoscale activity of sensory cortex.

## Methods

### Evicence for mesoscale models

Spontaneous reversals of perception exhibit statistical features that are universal in all well-studied cases and that tightly constrain the underlying neural architecture. First, spontaneous reversals reflect an interaction between two representations: a fast-changing representation of ‘dominant’ perceptual alternatives and a slowchanging representation of neural support (137–141). Second, perceptual alternatives interact through dynamic competition, which makes dominance durations depend sensitively on sensory input (71–73, 142, 143). Third, dominance durations are stochastic and their distribution exhibits a universal shape (approximating a Gamma-distribution) under all input conditions (54, 59, 60, 144, 145). This strongly suggests a birth-death process operating out-of-equilibrium, as this would guarantee the observed distribution shape under all conditions. (69, 70).

We investigated mesoscale models – with two or three alternative appearances – that comprehensively satisfy these constraints. The models are constructed from bistable ‘columns’ that ‘toggle’ spontaneously between low and high activity states (‘off’ and ‘on’). The kinetic rates of ‘toggling’ vary exponentially with external input (see below). This mimics ‘escapes’ from potential wells where kinetic rates generally depend exponentially on the potential difference (146). For attractor networks of spiking neurons, this exponential dependence has been confirmed by simulations and mean-field analyses (49, 70).

The two-alternative mesoscale model was specified fully by inverse modelling the reversal sequences of human observers for different combinations of two inputs, *I, I*′ (49). The empirical reversal statistics constrain the interaction strength within and between levels (excitatory, inhibitory, and destabilizing) as well as the population size at the lower level, as illustrated in **Suppl. Fig. 15** and **Fig. 16**. Only the upperlevel population size (here *N*_*r*_ = 10) was not constrained by perceptual evidence.

### Definition of mesoscale models

We idealize the spontaneous activity of a cortical column as being bistable and toggling with Poisson statistics between distinct ‘on’ and ‘off’ states (49). Toggling reflects ‘escapes’ from potential wells, so that the kinetic rates *v*^+^ and *v*^*−*^ depend exponentially and inversely on the mean-field potential difference Δ*u*(*t*)

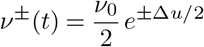

where *ν*_0_ is a baseline rate.

A population of *N* such bistable columns is described by the number *n*(*t*) of active columns or, equivalently, by the fractional activation *x*(*t*) = *n*(*t*)*/N*. The time-development of *n*(*t*) *∈* [0, …, *N*] is an inhomogeneous birth-death process and the probability *P*_*n*_(*t*) of *n* active units at time *t* is governed by the typical master equation of small-number discrete systems in statistical physics. When the potential difference Δ*u* is constant, the expected development ⟨*n*(*t*) ⟩ is simply exponential, with the steady-state increasing with input. A unique feature of birth-death processes is that the characteristic time-constant decreases with input (69).

The two-alternative model comprises four populations of toggling columns, two with *N*_*E*_ = 25 elements and two others with *N*_*R*_ = 10 elements, with fractional population activities *e*(*t*), *e*′(*t*), *r*(*t*), and *r*′(*t*), respectively. The kinetic rates 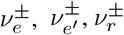 and 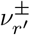 are uniform within each population and vary exponentially (see above) with the respective potential differences Δ*u*_*e*_, 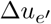, Δ*u*_*r*_ and 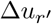.

Population activities (*e, e*′, *r, r*′) are coupled strongly to each other and to external input. Consistent with mean-field theory of spiking neurons, couplings are realized by linear dependencies between the activities of source populations and the potential differences governing the kinetics of target populations, as follows

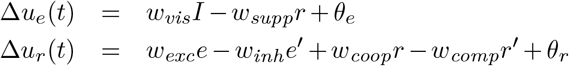

and symmetrically for 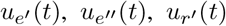 and 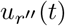. Model parameters *w*_*vis*_, *w*_*supp*_, *w*_*exc*_, *w*_*inh*_, *w*_*coop*_, and *w*_*comp*_ represent coupling strengths and model parameters *ν*_*e*_, *ν*_*r*_, *θ*_*e*_, and *θ*_*r*_ control toggling rates. External sensory input is represented by *I, I*′ *∈* [0, 1].

The three-alternative model comprises six populations of toggling columns, three with *N*_*E*_ = 25 elements and three others with *N*_*R*_ = 10 elements, with fractional population activities *e*(*t*), *e*′(*t*), *e*″(*t*), *r*(*t*), and *r*′(*t*), *r*″(*t*), respectively. The couplings are are

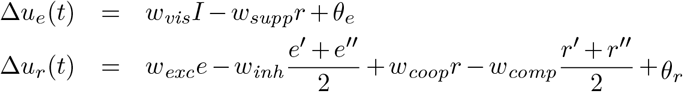

and symmetrically for *u*_*e*_ (*t*), *u*_*e*_ (*t*), *u*_*r*_ (*t*), and *u*_*r*_ (*t*). Note that upper-level interactions *w*_*coop*_ and *w*_*comp*_ were ‘diluted’ by a factor of 2 to account for the additional population.

All parameter values were determined by inverse modelling of reversal sequences experienced by human observers under variety of different input conditions *I, I*′, as described elsewhere (49). The fitted values were *ν*_*e*_ = 0.5128, *ν*_*r*_ = 55.556, *θ*_*e*_ = *−* 1.65, *θ*_*r*_ = *−* 4.94, *w*_*vis*_ = 1.78, *w*_*supp*_ = 2.34, *w*_*exc*_ = 120.1, *w*_*inh*_ = 32.1, *w*_*coop*_ = 15.21, and *w*_*comp*_ = 33.4.

### Macroscopic dynamics and dominance statistics

The simulated trajectories exhibit a macroscopic dynamic with alternating phases of ‘dominance’ and ‘reversals’. The criterion for ‘dominance’ of population *R* is defined as *r*(*t*) *≥ N*_*R*_ *−* 1 and *r*′(*t*) *≤ N*_*R*_*/*2, and analogously for *R*′. ‘Dominance periods’ are continuous periods of uninterrupted dominance. ‘Reversal periods’ are the transitions between dominance periods. The simulated trajectories comprised *O*(10^4^) cycles of dominance. Dominance durations are a stochastic variable, the statistics of which are given by distribution moments (duration, coefficient of variation, skewness) and by sequential correlations between successive durations (lagone-adjacent, lag-two-adjacent, and lag-three-adjacent). The mesoscale model is fully constrained by experimental observations with human observers of dominance statistics under different input conditions (*I, I*′).

The dominance statistics for different symmetric inputs *I* = *I*′ and for different scaling factors of horizontal and vertical _*e*_ interactions, *γ*_horiz_ and *γ*_vert_, are listed in **Tab. 1**. These values for unscaled interactions closely match the statistics of spontaneous reversals in the perception of binocular rivalry – with dominance durations measured in seconds – by a typical human observer (49).

**Table 1.** Dominance statistics (mean ± STD) of two-alternative models with different scaling factors, *γ*_horiz_ and *γ*_vert_, of interaction strengths and with different symmetric inputs *I* = *I*′.

| Model | γ<sub>vert</sub> | γ<sub>horiz</sub> | Input | Duration | Coeff. Var. | Skewness | CC Lag-1 | CC Lag-2 | CC Lag-3 |
|---|---|---|---|---|---|---|---|---|---|
| S | 1 | 1 | 0.00 | 4.0851 ± 0.0180 | 0.6682 ± 0.0026 | 2.7186 ± 0.0493 | 0.0059 ± 0.0092 | −0.0031 ± 0.0075 | 0.0078 ± 0.0042 |
| S | 1 | 1 | 0.25 | 3.0073 ± 0.0076 | 0.5576 ± 0.0025 | 2.9271 ± 0.0676 | 0.0001 ± 0.0031 | 0.0046 ± 0.0078 | 0.0015 ± 0.0046 |
| S | 1 | 1 | 0.50 | 2.4593 ± 0.0053 | 0.5006 ± 0.0030 | 2.9993 ± 0.0481 | 0.0304 ± 0.0036 | 0.0106 ± 0.0041 | 0.0030 ± 0.0024 |
| S | 1 | 1 | 0.75 | 1.7967 ± 0.0045 | 0.5269 ± 0.0017 | 2.3901 ± 0.0300 | 0.1351 ± 0.0026 | 0.0582 ± 0.0060 | 0.0205 ± 0.0060 |
| S | 1 | 1 | 1.00 | 1.0476 ± 0.0011 | 0.6629 ± 0.0011 | 1.7511 ± 0.0236 | 0.2883 ± 0.0058 | 0.1493 ± 0.0023 | 0.0761 ± 0.0061 |
| A | √2 | √2 | 0.75 | 1.8581 ± 0.0019 | 0.4674 ± 0.0023 | 2.7841 ± 0.0470 | 0.0690 ± 0.0050 | 0.0154 ± 0.0047 | 0.0068 ± 0.0064 |
| B | √2 | 1/√2 | 0.75 | 0.5860 ± 0.0010 | 0.6818 ± 0.0014 | 1.7375 ± 0.0219 | 0.2363 ± 0.0033 | 0.1295 ± 0.0037 | 0.0709 ± 0.0030 |
| C | 1/√2 | √2 | 0.75 | 15.0536 ± 0.1299 | 0.7496 ± 0.0087 | 2.4952 ± 0.1470 | 0.0015 ± 0.0158 | −0.0034 ± 0.0096 | −0.0078 ± 0.0128 |
| D | 1/√2 | 1/√2 | 0.75 | 1.7318 ± 0.0044 | 0.6745 ± 0.0032 | 1.8536 ± 0.0379 | 0.2316 ± 0.0056 | 0.1355 ± 0.0055 | 0.0834 ± 0.0044 |

### Rate of information gain

A birth-death process – defined as *N* bistable units with transition rates *ν*^*±*^ = *ν*_0_*e*^*±ws*^, where *ν*_0_ is a baseline rate and *w* a coupling constant – accumulates and retains information about input *s*, performing as a ‘leaky integrator’ with variable characteristic time (49). Specifically, the value of *s* may be inferred from fractional activity *x*(*t*) at time *t*, if coupling *w* and baseline rate *ν*_0_ are known. The inverse variance of the maximum likelihood estimate is given by the Fisher information

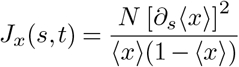

Its value grows with time, approaching *J*_*x*_ = *Nw*^2^*/* cosh^2^(*ws/*2) for *t →∞*. For small inputs *s ≃*0, the Fisher information increases monotonically as *J*_*x*_(*t*) *≈* (*Nw*^2^*/*4) tanh (*νt/*2). The upper bound of *J*_*x*_ *≤ N w*^2^*/*4 depends linearly on pool size *N*, but quadratically on coupling *w*. Thus, stronger coupling substantially improves the encoding of input *s*.

The rate at which Fisher information accrues depends on both population size *N* and baseline rate *ν*_0_. An initially inactive pool, with *x*(*t*) = 0, accumulates Fisher information at an initial rate of *∂*_*t*_*J*_*x*_| _*t*=0_ = *ν*_0_ (*Nw*^2^*/*4) *e*^(*ws/*2)^. Thus, any desired accumulation rate may be obtained by choosing appropriate values for *N* and *ν*_0_.

### Master equation and Markov model

For the twoalternative model, ‘microscopic’ trajectories of individual columns comprise 2 *× N*_*E*_ + 2 *× N*_*R*_ = 70 binary stochastic variables (‘toggling columns’). ‘Macroscopic’ trajectories of population activities comprise 4 integer stochastic variables, namely, the active columns in each of the four populations *e*(*t*), *e*′(*t*), *r*(*t*), *r*′(*t*). For the three-alternative model, ‘microscopic’ trajectories are 105-dimensional and ‘macroscopic’ trajectories 6-dimensional.

By enumerating and assigning indices *j, k*, … to the value quadruplets or sextuplets of macroscopic system ‘states’, one may equivalently generate trajectories from the Master equation of a first-order Markov model

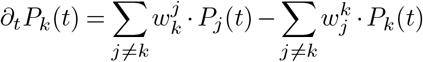

where rates 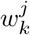(from *j* to *k*) and 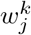 (from *k* to *j*) are given analytically by the model, for example

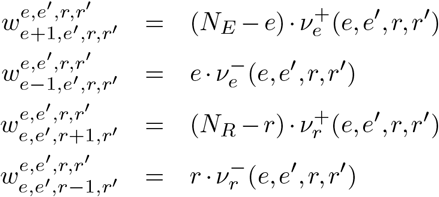

In practice, simulating state-space trajectories from a Markov model is numerically more challenging than simulating the birth-death dynamics of the mesoscale model.

The dynamical steady-state 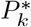 of a Markov dynamics is unique and defined by

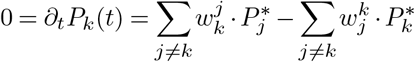

and may either be an equilibrium or a non-equilibrium steady-state (147).

The net steady-state flow between states *j* and *k* is given by

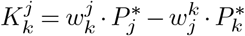

and vanishes for equilibrium steady-states.

### Inferring dynamic Hamiltonian functions

A dynamic Hamiltonian function for the two-alternative model defines the joint probability of ‘on-off toggles’ in terms of population activities *e, e*′ *∈* {0, 1, …, *N*_*e*_*}* and *r, r*′ *∈ {*0, 1, …, *N*_*r*_*}* and on up to eleven inferred Lagrange multipliers, 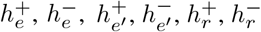, *K*_*coop*_, *K*_*comp*_, *K*_*exc*_, *K*_*inh*_, *K*_*supp*_.

Multipliers 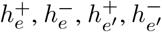 determine the basic toggling rate of lower-level populations *E, E*′, including the effect of the sensory inputs, *I, I*′. For symmetric input *I* = *I*′, only two multipliers 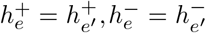 are needed. Similarly, the multipliers 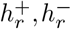 set the basic toggling rate of upper-level populations *R, R*′.

Multipliers *K*_*exc*_, *K*_*inh*_, *K*_*coop*_, *K*_*comp*_, *K*_*supp*_ represent the feedforward excitation and inhibition from lowerto upper-level populations (*K*_*exc*_, *K*_*inh*_), the cooperative and competitive interactions between upper-level populations (*K*_*coop*_, *K*_*comp*_), and the feedback suppression from upper- to lower-level populations (*K*_*supp*_).

The toggle counts 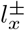 in the dynamic Hamiltonian function can be realized either with binary random variables 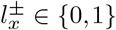 or with uniformly distributed random variables 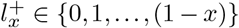 and 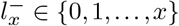, to better accommodate bursts of toggles. In the present work on mesoscopic models, the time interval *dt* = 0.0001 s was so short that both approaches yielded equivalent results.

Starting from long simulated trajectories (9.6 *×* 10^4^ seconds, 9.6 *×* 10^8^ time intervals) in duration, we used Markov Chain Monte Carlo to infer the values of Lagrange multipliers. To ensure convergence, 2.5 *×* 10^5^ iterations were generated. The resulting values are listed in **Tab. 2**.

**Table 2.** Lagrange multipliers (mean ± STD) of dynamic Hamiltonian function, for two-alternative models with different scaling factors, *γ*_horiz_ and *γ*_vert_, and different sensory inputs, *I* and *I*′.

| Model | S | S | A | B | C | D |
| γ<sub>vert</sub> | 1 | 1 | √2 | √2 | 1/√2 | 1/√2 |
| γ<sub>horiz</sub> | 1 | 1 | √2 | 1/√2 | √2 | 1/√2 |
| Input *I* | 0.25 | 0.75 | 0.75 | 0.75 | 0.75 | 0.75 |
| Input *I*′ | 0.25 | 0.75 | 0.75 | 0.75 | 0.75 | 0.75 |
|---|---|---|---|---|---|---|
| *h*<sub>*e*</sub><sup>+</sup> | −11.26 ± 0.17 | −10.81 ± 0.14 | −10.84 ± 0.14 | −10.83 ± 0.14 | −10.83 ± 0.15 | −10.81 ± 0.14 |
| *h*<sub>*e*</sub><sup>−</sup> | −10.08 ± 0.23 | −10.55 ± 0.21 | −10.49 ± 0.23 | −10.51 ± 0.27 | −10.50 ± 0.17 | −10.53 ± 0.19 |
| *K*<sub>*exc*</sub> | 2.68 ± 0.14 | 2.700 ± 0.106 | 3.65 ± 0.16 | 3.786 ± 0.095 | 1.87 ± 0.20 | 1.865 ± 0.071 |
| *K*<sub>*inh*</sub> | 0.688 ± 0.067 | 0.727 ± 0.61 | 1.01 ± 0.10 | 1.01 ± 0.050 | 0.50 ± 0.14 | 0.502 ± 0.041 |
| *K*<sub>*supp*</sub> | 0.120 ± 0.026 | 0.122 ± 0.21 | 0.162 ± 0.024 | 0.164 ± 0.028 | 0.080 ± 0.019 | 0.085 ± 0.019 |
| *h*<sub>*r*</sub><sup>+</sup> | −8.41 ± 0.51 | −8.60 ± 0.45 | −8.23 ± 0.67 | −8.65 ± 0.24 | −8.26 ± 1.56 | −8.47 ± 0.41 |
| *h*<sub>*r*</sub><sup>−</sup> | −3.48 ± 0.54 | −3.37 ± 0.46 | −3.62 ± 0.75 | −3.39 ± 0.22 | −3.75 ± 1.67 | −3.45 ± 0.41 |
| *K*<sub>*coop*</sub> | 0.781 ± 0.060 | 0.821 ± 0.049 | 1.125 ± 0.085 | 0.577 ± 0.023 | 1.10 ± 0.20 | 0.561 ± 0.033 |
| *K*<sub>*comp*</sub> | 1.88 ± 0.11 | 1.875 ± 0.084 | 2.55 ± 12 | 1.327 ± 0.40 | 2.60 ± 0.29 | 1.300 ± 0.058 |

### Analytical relation between master equation and Hamiltonian function

#### Similarity of trajectories

To quantify the similarity or dissimilarity of the stochastic trajectories generated by different simulations or systems, one can compare the spaces sampled by stochastic fluctuations over time. A well-established quantitative measure is the difference between the roots of covariance matrices, relative to the total variance (‘Hess overlap’)(148).

Specifically, the covariance matrices *A* and *B* of two trajectories are used to obtain the matrix difference *d*(*A, B*) and the matrix overlap *s*(*A, B*):

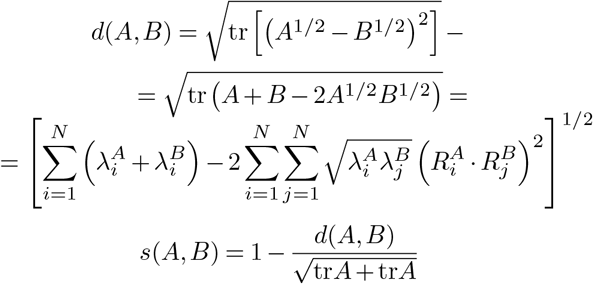

where the square root of the covariance matrix *A* is defined as

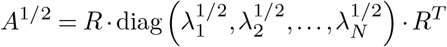

and *R*_*i*_ is the *i*th column of rotation matrix *R*. The overlap *s*(*A, B*) is unity if matrices *A* and *B* and sampled subspaces are identical and zero when the sampled subspaces are orthogonal.

*Mesoscopic* trajectories of population activities (*e, e*′, *r, r*′) are four-dimensional, yielding (4 *×* 4) covariance matrices. The *microscopic* trajectories of bistable column activities are 70-dimensional (2 *×* 25 lower-level columns and 2 *×* 10 upper-level columns), so that covariance matrices are (70 *×* 70).

Trajectories of two-alternative models were simulated 96000 s and stepsize *dt* = 0.1 ms. The Hess overlap between the first and second trajectory halves was *≥* 0.995 for all macroscopic trajectories and *≥* 0.98 for all microscopic trajectories, confirming convergence for all simulated models and input conditions (**Tab. 3**).

**Table 3.** Hess overlap between trajectories from the same model, including the standard model S for five different inputs *I* = *I*′, and from modified models A through D for one input *I* = *I*′ = 0.75.

| Model | γ<sub>vert</sub> | γ<sub>horiz</sub> | Input<br>*I* = *I*′ | Mesoscopic overlap | Microscopic overlap |
|---|---|---|---|---|---|
| S | 1 | 1 | 0.00 | 0.9989 | 0.9859 |
| S | 1 | 1 | 0.25 | 0.9979 | 0.9858 |
| S | 1 | 1 | 0.50 | 0.9982 | 0.9852 |
| S | 1 | 1 | 0.75 | 0.9981 | 0.9834 |
| S | 1 | 1 | 1.00 | 0.9991 | 0.9831 |
| A | √2 | √2 | 0.75 | 0.9982 | 0.9865 |
| B | √2 | 1/√2 | 0.75 | 0.9986 | 0.9848 |
| C | 1√2 | √2 | 0.75 | 0.9966 | 0.9818 |
| D | 1/√2 | 1/√2 | 0.75 | 0.9970 | 0.9809 |

To assess the similarity of macroscopic trajectories from different two-alternative models and different dynamic Hamiltonian functions, we computed the Hess overlap of trajectory pairs from original models, pairs from dynamic Hamiltionians, and mixed pairs from both methods. The results are shown in **Suppl. Fig. 9b**.

#### Observed flow

We can obtain trajectory flow and transition rates from a temporal sequence of space vector locations (*E, E*′, *R, R*′) of sufficient length *N*. For simplicity, we assume that the sequence is cyclical and that the first location succeeds the last location. From the sequence, we can obtain the fractional occurrence *f*_*k*_ of location *k*

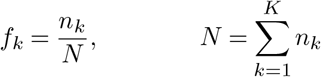

where *n*_*k*_ is the observed number of visits to location *k*. In addition, we can obtain the fraction 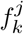 of transitions between distinct locations, the total number of such transitions *N*_*trans*_, and the overall rate *r*_*trans*_ of transitions:

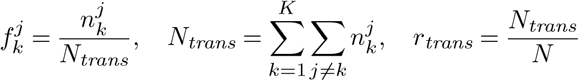

where 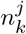 is the observed number of transitions from location of origin *j* to location of destination *k* ≠ *j*. The simulated trajectories comprised *O*(10^6^) transitions between distinct locations in state space. As any transition is *from* one location *to* another location, row and column sums in the matrices 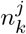 and 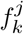 are equal

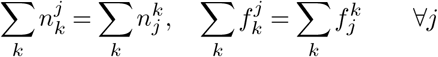

Taking the observed occurrence fractions *f*_*k*_ of states *k* as estimates for steady-state probabilities

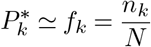

we can estimate transition rates 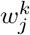 in terms of the flow rates implied by the observed transition counts

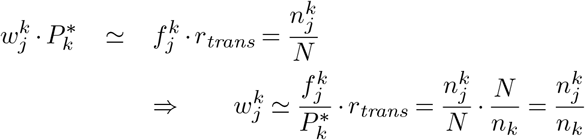

Note that these estimates are guaranteed to satisfy the steadystate condition *∀k*

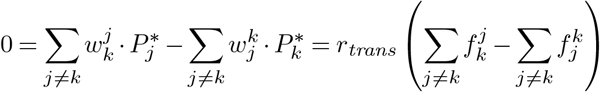

#### Entropy production

The commonly used expression for entropy production in non-equilibrium systems (147, 149) is the weighted sum of imbalanced flows

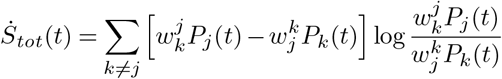

where the sum is taken over all *unordered* pairs *k* ≠ *j*. This total entropy production can be divided into contributions of the ‘system’ and the ‘medium’, 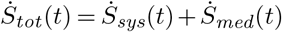, with

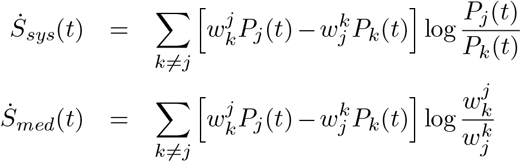

The system contribution is the time derivative of entropy 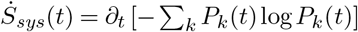 and vanishes for a time-invariant steady-state distribution 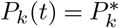. Here we use 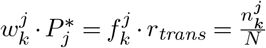 to obtain

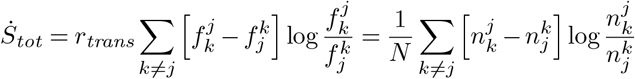

Each unordered pair *k ≠ j* contributes

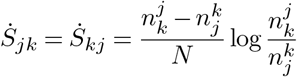

to the total entropy production.

When no transitions are observed for a subset of ordered pairs *k →j*, the associated transition flows vanish and the entropy production becomes arbitrarily large (147). For this reason, entropy production does not provide a stable measure for ‘distance-from-equilibrium’.

As a rough approximation, we compute entropy production for all transitions observed in both directions (*k →j* and *j →k*), disregarding transitions observed only in one direction. These ‘bidirectional’ transitions included about 85% of the *O*(10^6^) transitions in a simulated trajectory.

#### Net flow and ‘bad actor’ flow

As alternative measures for distance from equilibrium, we obtained from simulated trajectories the observed forward, reverse, and net flows for every ordered pair of states *j, k*

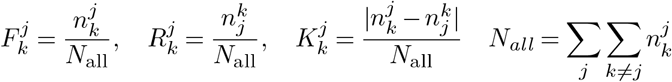

as well as the summed totals

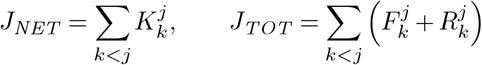

If all net flows vanish, detailed balance is maintained and the dynamics are near equilibrium. If not, detailed balance is broken, and the dynamics are non-equilibrium. Thus, a convenient measure for distance from equilibrium is the net flow as a fraction of total flow:

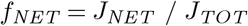

This measure is unity if the flows between all state pairs are maximally unbalanced (unidirectional) and zero if all flows are exactly balanced.

The system in question is evidently not at equilibrium, as trajectories pursue a stochastic limit cycle in state space, as made evident by projecting onto the plane defined by differential activities, *e − e*′ and *r − r*′. Accordingly, we can define the predominant rotational sense in terms of

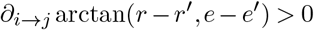

With respect to this predominant rotation, we can classify all transitions *i →j* as either ‘good actor’, ‘bad actor’, or ‘neutral actor’ (with, against, or normal to the rotation, respectively). For purposes of comparison, we can analogously classify transitions with respect to other, orthogonal axes, such as *e* + *e*′ or *r* + *r*′, where we do not expect any predominant direction of motion.

With respect to these reference directions, we obtained from simulated trajectories the observed ‘good’ flows (with the direction) and ‘bad’ flows (against the direction) as

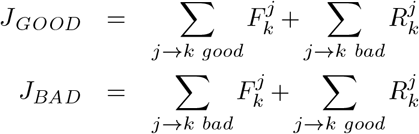

To quantify distance from equilibrium, we can use the ratio of good to bad flows, or the difference between good and bad flows as a fraction of total flow:

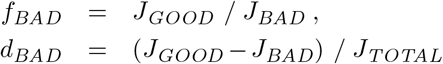

#### Trajectory caliber

To characterize the non-equilibrium dynamics in more detail, we parsed simulated trajectories into non-overlapping periods of ‘dominance’ (as defined above) and ‘reversals’ (the intervening intervals), with each trajectory segment comprising of a sequence of distinct transitions. To quantify the variability of these segments, we computed path entropy (‘caliber’) separately for ‘dominance’ and ‘reversal’ segments. Given an ensemble of *O*(10^4^) trajectory ^*j*^segments from one simulated trajectory, we established the list of *O*(10) unique sequence types Γ, differentiating the total count of increments and decrements in the four state variables:

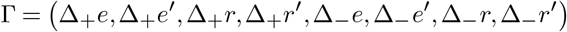

Note that this focus on the variability of *counts* disregards variability in the *order* or the *timing* of individual increments and decrements. For this reason, our path entropy values are likely to systematically *underestimate* the true value. Given a list of sequence types Γ, we obtained the probability of each type and computed path entropies as

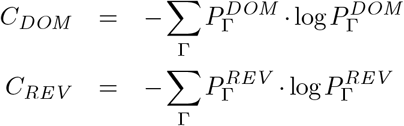

distinguishing between dominance and reversal sequences. For purposes of comparison, we also established a list of sequence lengths (total number of increments and decrements) and assessed the variability of sequence length in terms of length entropy.

#### Simulation of spiking neurons

To generate predictions for spiking neurons, we simulated the three-alternative model in terms of the binary states (‘on’ or ‘off’) of the 105 columns from six populations (3 *×* 25 + 3 *×* 10). To ensure that toggling rates resemble those observed in early sensory cortex, we elevated baseline rates twenty-fold, to *ν*_*e*_ = 10.256 and *ν*_*r*_ = 1111.12. The resulting mean toggling rates were 2.9 Hz and 3.5 Hz for columns in lower- and upper-level populations, respectively. Specifically, we simulated 100 trials of 6 s duration.

As described in *Section 2*.*5*, the three sensory inputs, *I*(*t*), *I*′(*t*), *I*″(*t*), were modulated during each trial. While the baseline value of all three inputs was 0.3, input *I* was elevated to 0.6 and input *I*′ reduced to 0.2 from 2 s to 4 s after the start of each trial.

In a second step, we randomly selected two columns from each of the upper-level populations *r, r*′, *r*″ and lower-level populations *e, e*′, *e*″ (12 columns in total), and generated Poisson spike trains from the toggling time-course of each column. Specifically, we generated spike trains as an inhomogeneous Poisson process with a rate of either 10 Hz or 50 Hz, depending on whether the column was ‘off’ or ‘on’. For each selected column, 10 independent spike trains were generated in this way, providing 120 spike trains in total.

#### Correlations of spiking activity

Correlations of spiking activity were analyzed with the conventional methods (101), summarized below.

Spike-trains are discrete binary series at millisecond resolution, 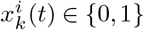, with neuron *k ∈ {*1, 2}, trial *i* (1 *≤ i ≤ M*), and time *t* (1 *≤ t ≤ T*). The spike-counts for trial *i* and the post-stimulus-time-histogram (PSTH) over all trials are

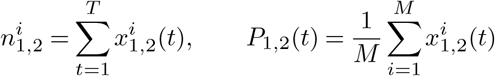

‘Noise correlation’ refers to shared trial-by-trial fluctuations in the spike counts of two neurons under the same stimulus condition. When the spike counts 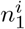 and 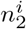 are observed over multiple trials *i*, the noise correlation *r*_*SC*_ is

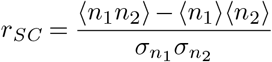

where ⟨⟩ is the average over trials and *σ*_*n*_ is the standard deviation of spike counts over trials.

Coincident spikes were counted by auto- or cross-correlation functions of spike trains or of PSTHs

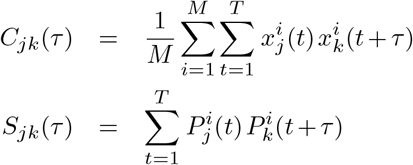

The auto- or cross-correlogram *CCG*(*τ*) of spike times is

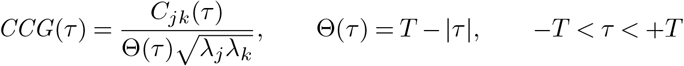

where *λ*_*j,k*_ are the mean firing rates (spikes per second) and Θ(*τ*) is a triangular function representing the overlap between spike trains of duration *T*.

The area under the auto- or cross-correlations integrated from *−T* to +*T* (after subtracting the shift-predictor) is

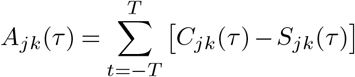

and the ratio of areas between cross- and auto-correlations is

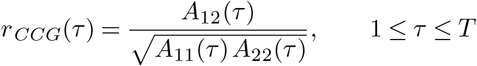

This measure compares the integrals of the central regions of CCG and AAG and asymptotes to the value of *r*_*SC*_ for *τ →T* :

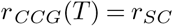

#### Fisher information

To compute the information about alternative inputs encoded by population activity, we represented the 60 upper-level spike trains as spike rates (per second) during successive 100 ms intervals. For each interval, we collected the observed vectors 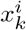 and opponent vectors 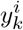 (in which the spike rates from populations *r* and *r*′ were swapped) from all trials, where *k* indexes *K* = 60 neurons and *i* indexes *N* = 100 trials, and computed averages and covariances over trials:

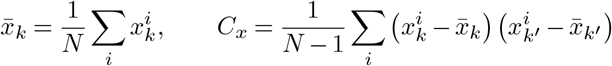

The Fisher information about alternative inputs for each 100 ms interval was then

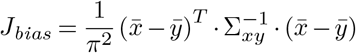

where 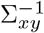 is the inverse of the average covariance Σ_*xy*_ = (*C*_*x*_ + *C*_*y*_) */*2. Applying the standard correction for bias due to small numbers of trials and neurons provides the corrected value

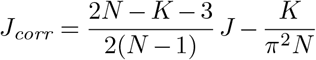

To obtain Fisher information from shuffled time series, we permuted the spike rates of each neuron within five consecutive 100 ms intervals.

#### Arrow-of-time irreversibility

Arrow-of-time irreversibility (5, 150, 151) quantifies the difference between time-series in positive (forward) and negative (reversed) time and provides a physics-based alternative for information-theoretical measures of causal direction and influence (152). Given two time-series *x*(*t*) and *y*(*t*), we denote *ρ*_*x→y*_(*τ*) and *ρ*_*y→x*_(*τ*) the lagged correlations

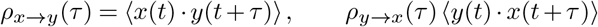

where *τ ≥* 0 is a positive lag. These lag-dependent correlations can be converted to lag-dependent mutual information

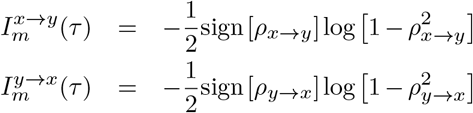

where the sign of the correlation is retained. The lag-dependent irreversibility is the squared difference,

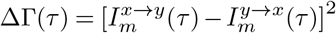

and the area under this curve, *A*_ΔΓ_ = ∫ΔΓ(*τ*)*dτ*, measures total irreversibility and causal influence.

Statistical significance is assessed by comparing the area *A*_ΔΓ_ of the observed time-series with a null-distribution obtained from randomly shifted time-series.

If there is significant irreversibility, the putative direction and type of causal influence is revealed by delayed peak correlations. For example, a positive peak of *ρ*_*x→ y*_ (*τ*) for some value *τ >* 0 indicates a facilitatory influence from *x*(*t*) to *y*(*t*). Similarly, a negative peak for some value *τ >* 0 would indicate a suppressive influence in the same direction.

## Acknowledgments

JB thanks Andrew Parker and Maurizio Mattia for helpful comments and the Otto-von-Guericke University for support.

## Supplementary material

### Trajectory comparison

To assess dynamical similarity, the original and Hamiltonian trajectories were compared in several qualitative and quantitative ways. Note that the correspondence between original and Hamiltonian trajectories is not expected to be exact, because the Hamiltonian is merely a linear approximation of the original dynamic couplings.

Direct visual comparisons of the trajectory ensembles generated by the original and Hamiltonian models, respectively, are particularly illuminating. This is possible in terms of either average trajectory flows, *∂*_*t*_ (*⟨ e⟩, ⟨ e*′*⟩, ⟨ r⟩, ⟨ r*′*⟩*), or average trajectory dispersions, *∂*_*t*_ (*⟨ e*^*2*^ *⟩ − ⟨e ⟩*^*2*^, *⟨e* ^*’2*^*⟩ − ⟨e*′ *⟩*^*2*^, *⟨r* ^2^*⟩ − ⟨r⟩* ^*2*^, *⟨r* ^*’2*^ *⟩ − ⟨r* ′ *⟩*^*2*^), which can be visualized in three-dimensional subspaces such as (*e − e*′, *r − r*′, *e* + *e*′) or (*e − e*′, *r − r*′, *r* + *r*′) (please see Methods for details).

Average flows from the 128 most visited locations with categorical dominance (*i*.*e*., *r − r*′ = *±*1) are compared in **Suppl. Fig. =9a** and average dispersions in **Suppl. Fig. 10**. It is evident that the correspondence is close and that any residual differences are small. Correspondence is comparably good for average flows and dispersions from other locations (for example, locations without dominance, *r − r*′ = 0, results not shown).

As mentioned, our theoretical expectation was that, however close, the correspondence would not be exact. More quantitative comparisons such as reversal statistics and Hess overlap confirmed this expectation. While the reversal statistics – duration *T*_dom_, coefficient of variation *C*_*V*_, skewness *γ*, lagged correlations *cc*_1_ – of the original and Hamiltonian dynamics were very similar and differed by less than 0.5%, the residual differences were systematic and became significant for sufficiently long trajectories with *O*(10^4^) reversals. Similarly, the Hess overlap between original and Hamiltonian trajectories was high with a value of 0.9900, but was not quite as high as the overlap between two independently generated original trajectories (0.9981) or two independent Hamiltonian trajectories (0.9987). In conclusion, the residual differences between original and Hamiltonian dynamics were small but systematic.

To assess trajectory similarity more comprehensively, we extended the investigation to mesoscale models that were modified in terms of either input strength *I* = *I*′ or interaction strength *w*_*coop*_, *w*_*comp*_, *w*_*exc*_, *w*_*inh*_, or *w*_*supp*_ and that therefore generated altered dynamics (see Methods for details). For each modified model with altered dynamics, we obtained both original trajectories and the corresponding Hamiltonian trajectories.

Having multiple pairs original and Hamiltonian trajectories allowed us to compare trajectory similarity and dissimilarity within and between pairs. Representative results are illustrated in **Suppl. Fig. 9b** for five models with different interaction strengths – dubbed S, A, B, C, D (see Methods for details). In terms of the Hess overlap measure, trajectory pairs from the same model are consistently similar, whereas trajectory pairs from different models are consistently more dissimilar, confirming the usefulness of this measure. More importantly, the pattern of pairwise Hess overlap was similar for original and Hamiltonian trajectories (middle panel), for pairs of original trajectories (upper panel), and for pairs of Hamiltonian trajectories (lower panel). This confirms that the dynamics generated by the original and the Hamiltonian models were nearly interchangeable.

The reversal statistics of original and Hamiltonian trajectories are compared in **Suppl. Fig. 9c**. Duration *T*_dom_, coefficient of variation *C*_*V*_, and skewness *γ* were consistently similar and differed by less than 0.5%, 1%, 1%, 1%, and 4%, respectively, for Models S, A, B, C, and D. As mentioned, the residual differences were systematic and became highly significant for sufficiently long trajectories with *O*(10^4^) reversals.

The largest residual differences between original and Hamiltonian trajectories were obtained for model D, as evident from both the reversal statistics and the Hess overlap. The latter value was 0.979 for model D, well below the overlap of two independent original trajectories (0.997) or two independent Hamiltonian trajectories (0.998). Interestingly, this larger discrepancy appeared to be more attributable to trajectory dispersion than to trajectory flow, as illustrated in **Suppl. Fig. 11**.

#### Further posterior distributions

To establish generality, we inferred Lagrange multiplier values for also trajectories generated by models A, B, C, and D in which the strength vertical and/or horizontal interactions had been scaled compared to the original model S. The resulting posterior distributions of multiplier values are illustrated in **Suppl. Fig. 12-13**. These figures also show the original multiplier values of model S (solid white crosshairs), as well as the values predicted by linearity (dashed white crosshairs). The excellent correspondence between the inferred posterior distributions and the predictions shows that multiplier values change approximately linearly with the strength of the underlying interactions.

#### Trajectory ‘inertia’ at diffusion limit

In the diffusion limit with infinite populations, dynamic trajectories become deterministic and seem to have ‘inertia’ even in the presence of external noise. **Suppl. Fig. 14** illustrates this ‘inertia’ with trajectory segments of the three-alternative model subjected to moderate noise and strong noise. Trajectories occasionally invert cycle direction only for strong noise.

### Constraints from perceptual evidence

The mesoscale model was constrained by perceptual evidence comprising four reversal statistics (duration, variance, skewness, lagged correlation) and two input strengths. To visualize these constraints, we illustrate selected cuts through this 6D space. **Suppl. Fig. 15** shows reversal statistics (for input *I* = *I*′ = 0.5) changing with horizontal and vertical interaction strength, revealing how the former constrain the latter. **Suppl. Fig. 16** shows reversal statistics changing with population size *N*_*E*_ and symmetric input *I* = *I*′, revealing that skewness and lagged correlations for strong inputs constrain the value of *N*_*E*_ *≈* 25.

**Fig. 9.**
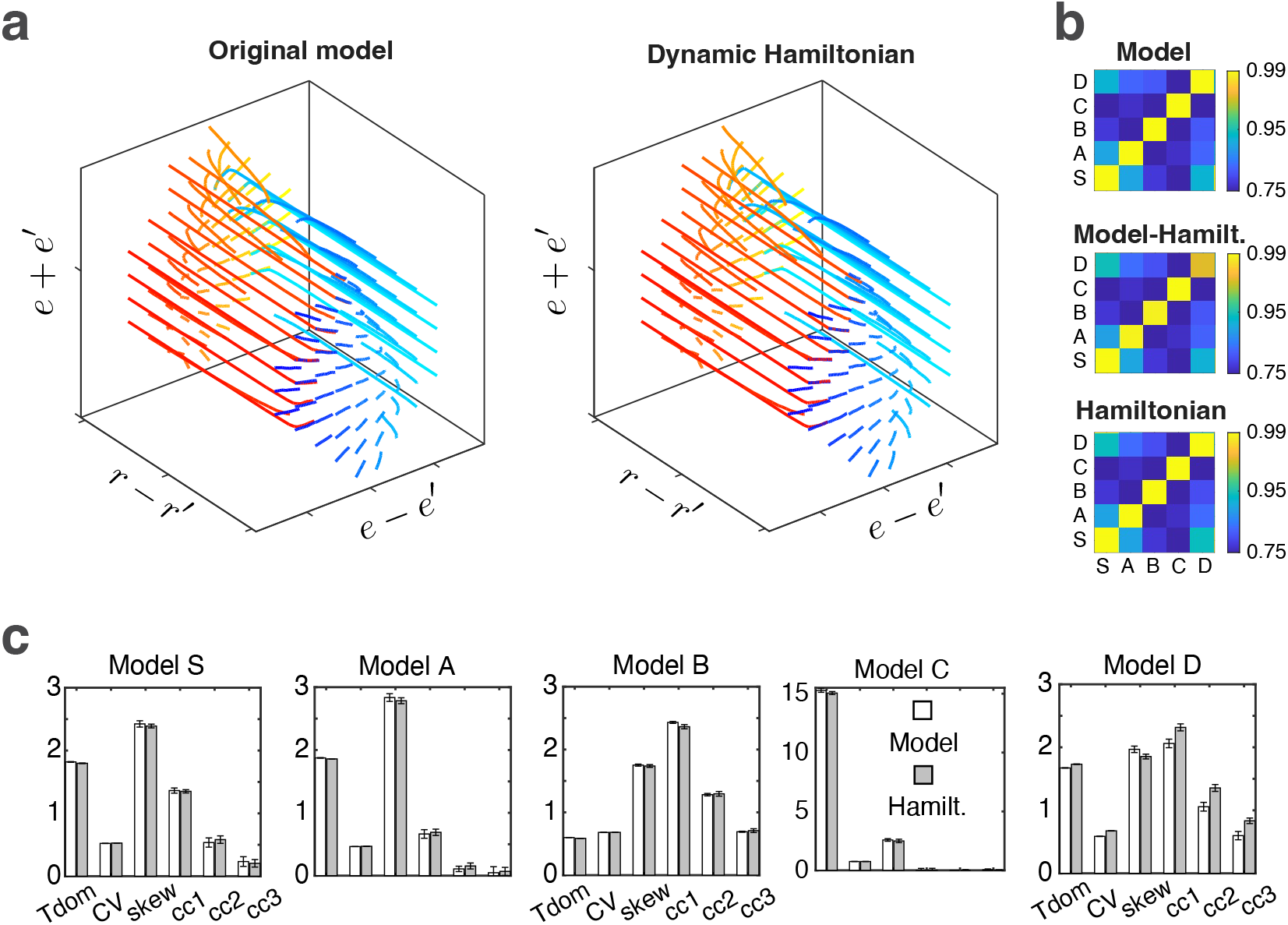
Asymptotic equivalence of model and Hamiltonian dynamics. **(a)** Average trajectory flow from selected initial locations in three-dimensional subspace (*e − e*′, *r − r*′, *e* + *e*′). Iterative simulations of the original dynamic (left) and of the corresponding minimal Hamiltonian dynamic (right) produce nearly identical trajectory flows. All trajectory durations were 0.2 s. Note slower flow in (*E, E*′) planes and faster flow in *R − R*′ direction. **(b)** Pairwise similarity of the trajectories generated by five different dynamics (models S, A, B, C, D). Similarity is measured in terms of the Hess overlap of the state spaces sampled by trajectory fluctuations. This overlap is comparable for pairs of CC trajectories (left), pairs of Hamiltonian trajectories (right), and mixed pairs of model and Hamiltonian trajectories (middle). **(c)** Statistics of perceptual reversals generated by five dissimilar model dynamics (light columns) and the corresponding Hamiltonian dynamics (grey columns). Corresponding values of dominance duration, *T*_*dom*_, coefficient of variation, *CV*, skewness, *γ/CV*, and sequential correlation, *cc*1, *cc*2, *cc*3, are nearly identical. Please note that sequential correlation values were scaled 10-fold for clarity.

**Fig. 10.**
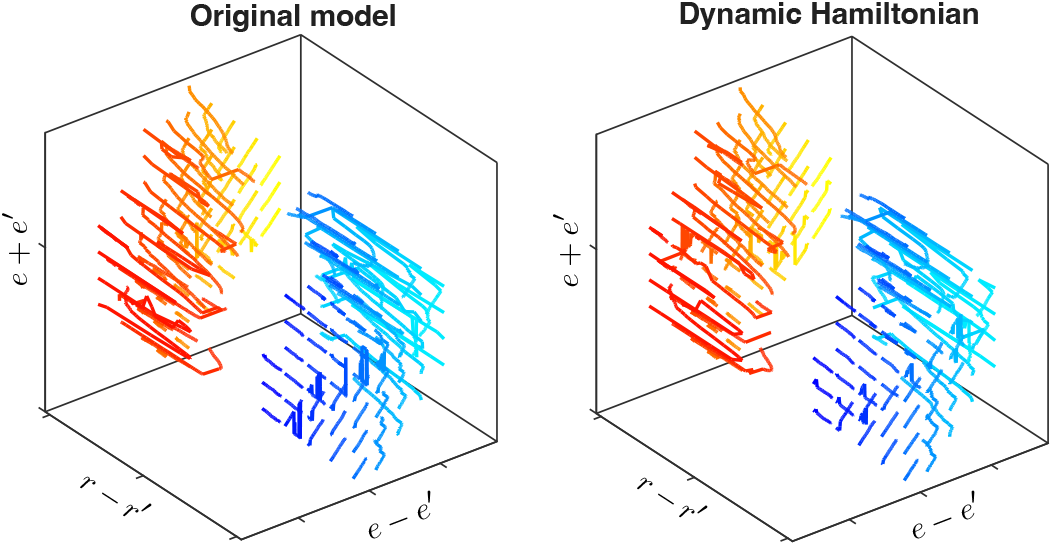
Asymptotic equivalence of original model and Hamiltonian dynamics. Average trajectory dispersion from selected initial locations in three-dimensional subspace (*e − e*′, *r − r*′, *e* + *e*′). Iterative simulations of the original CC dynamic (left) and of the corresponding minimal Hamiltonian dynamic (right) produce nearly identical trajectory dispersions. All trajectory durations were 0.2 s.

**Fig. 11.**
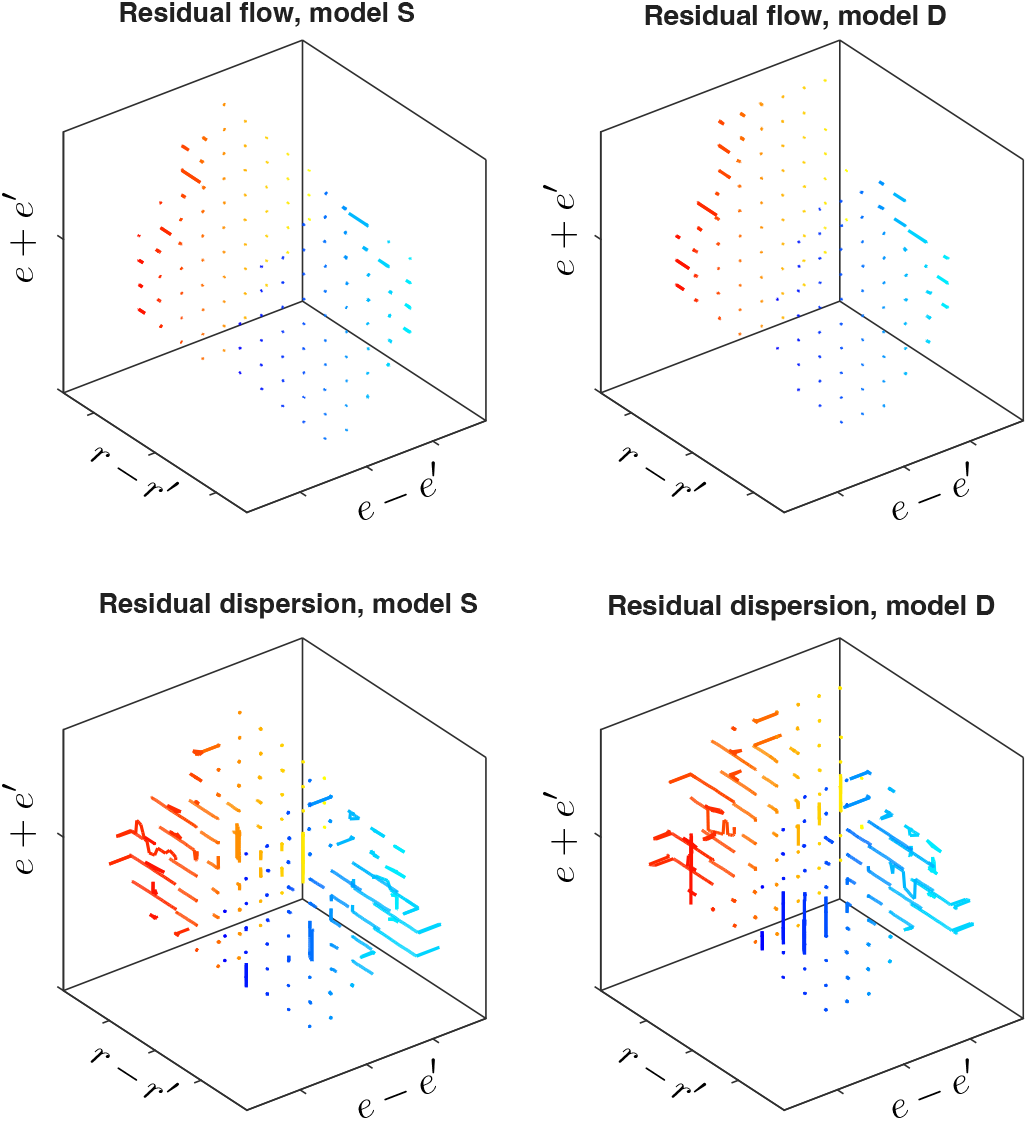
Residual differences between original model and Hamiltonian dynamics. Residual difference in average trajectory flow (top row) and average trajectory dispersion (bottom row), for Model S (left) and Model D (right). The main difference between Models S and D concerns dispersion during reversals.

**Fig. 12.**
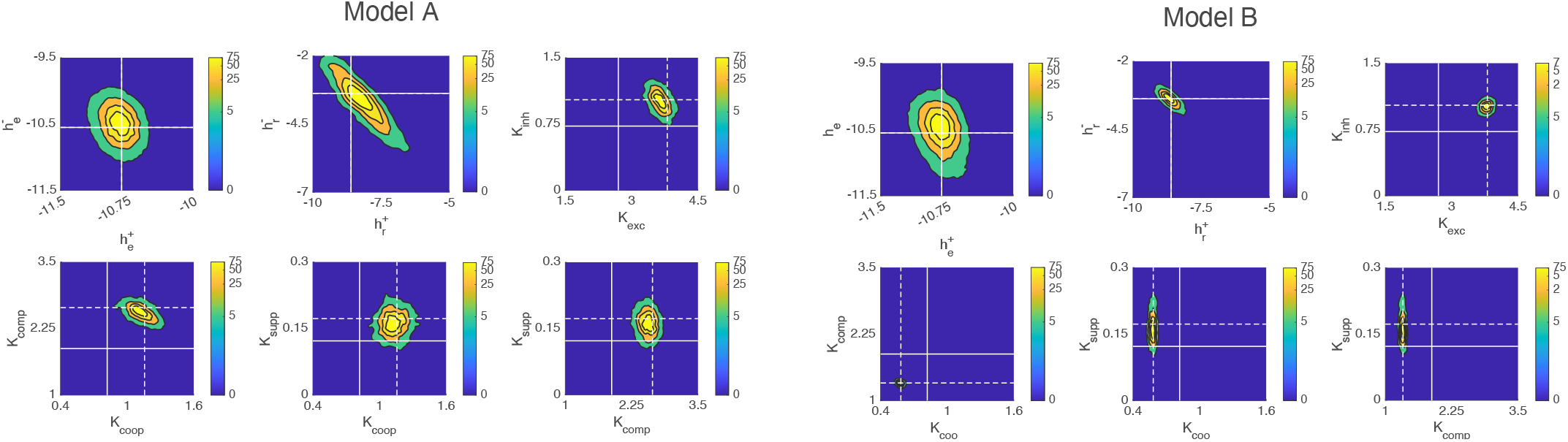
Posterior distribution of Lagrange multipliers for Models B and C. Joint posterior likelihood of Lagrange multipliers 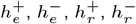, *K*_*coop*_, *K*_*comp*_, *K*_*exc*_, *K*_*inh*_, and *K*_*supp*_, fitted to a long trajectory *e, e*′, *r, r*′ of the original CC model C, under the constraint of Maximum Likelihood. Color contours represent the cumulative posterior probability over the regions *external* to each contour.

**Fig. 13.**
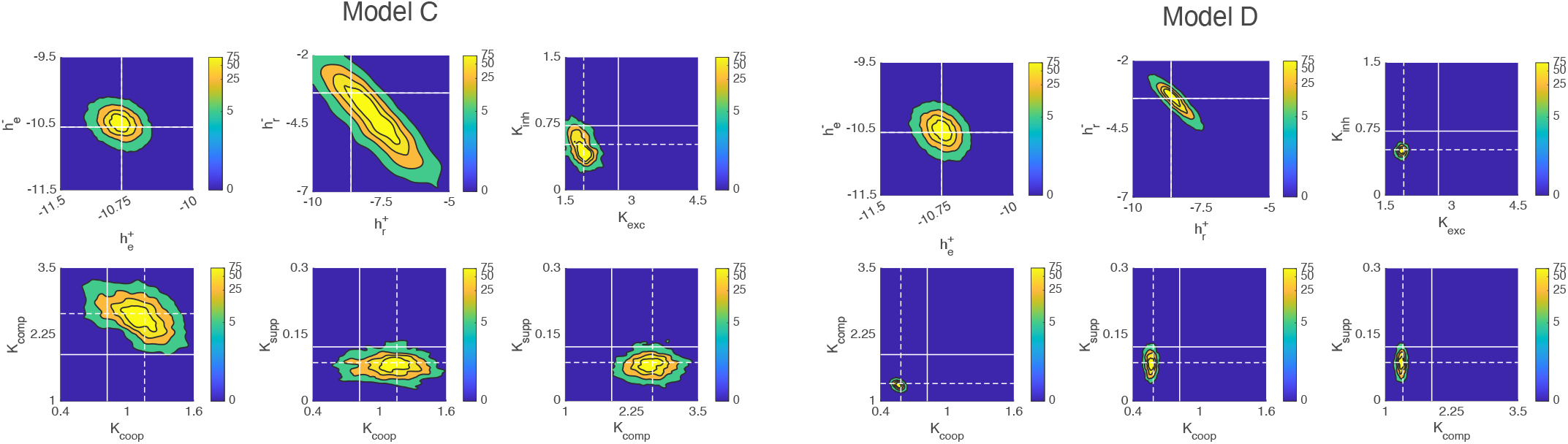
Posterior distribution of Lagrange multipliers for Models B and C. Joint posterior likelihood of Lagrange multipliers 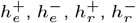, *K*_*coop*_, *K*_*comp*_, *K*_*exc*_, *K*_*inh*_, and *K*_*supp*_, fitted to a long trajectory *e, e*′, *r, r* ′ of the original CC model C, under the constraint of Maximum Likelihood. Color contours represent the cumulative posterior probability over the regions *external* to each contour.

**Fig. 14.**
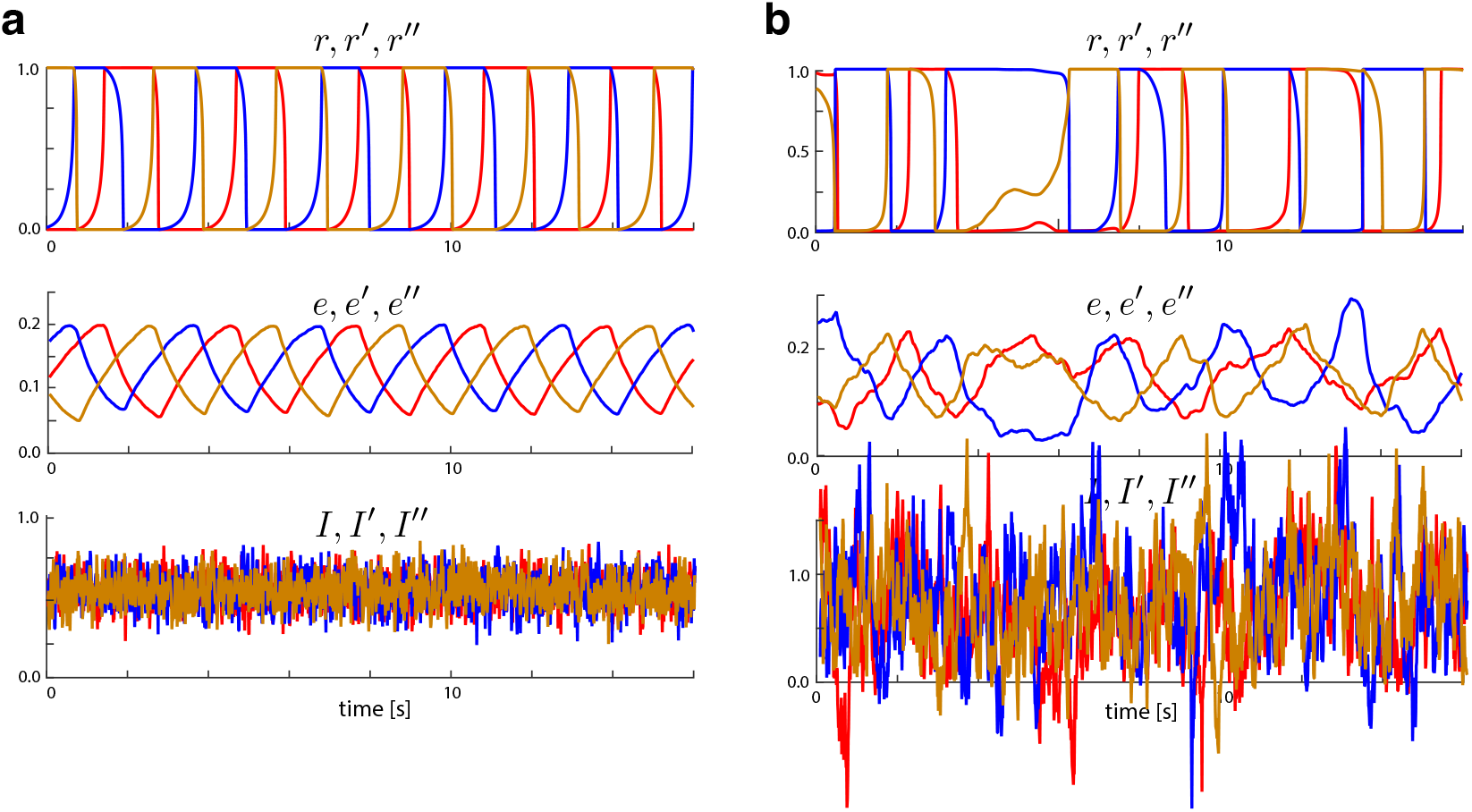
Deterministic dynamic at ‘diffusion limit’. **(a)** At the ‘diffusion limit’, *N*_*e*_, *N*_*r*_ *→∞*, trajectories are deterministic and follow a limit cycle, with the cycle direction determined by initial conditions. This ‘trajectory inertia’ persists even with variable sensory input (here *µ*_*I*_ = 0.5 and *σ*_*I*_ = 0.1). **(b)** Inverting the cycle direction requires strong external forcing (here *µ*_*I*_ = 0.5 and *σ*_*I*_ = 0.5), which exceeds the normal dynamic range of input, [0, 1].

**Fig. 15.**
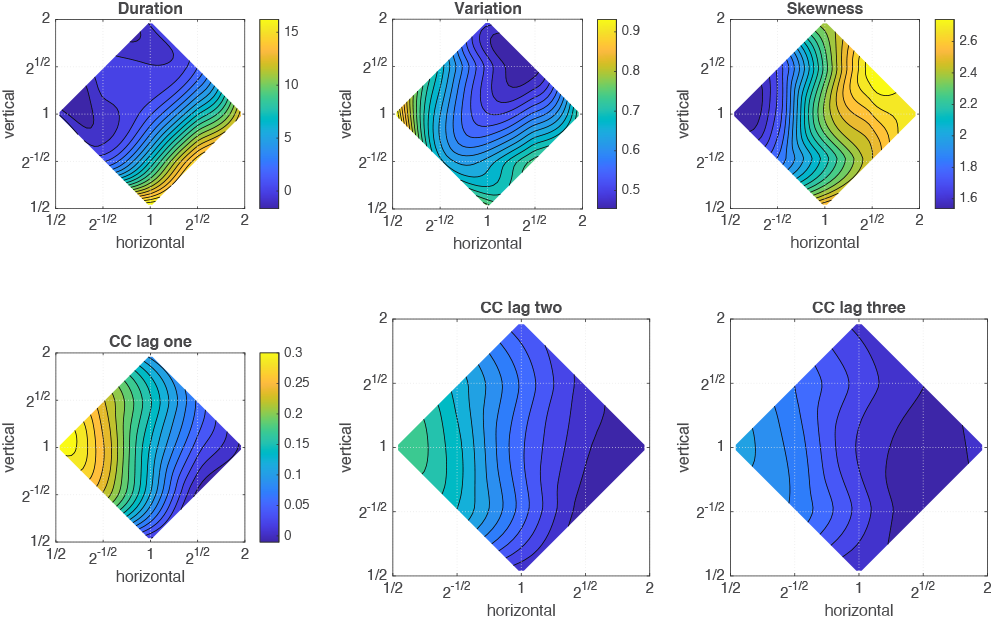
Perception constrains interaction strength. The values of six reversal statistics – dominance duration, coefficient of variation, skewness, sequential correlations for lag one, two, and three – change differentially with the strength of upper-level interactions (‘horizontal’) and of inter-level interactions (‘vertical’).

**Fig. 16.**
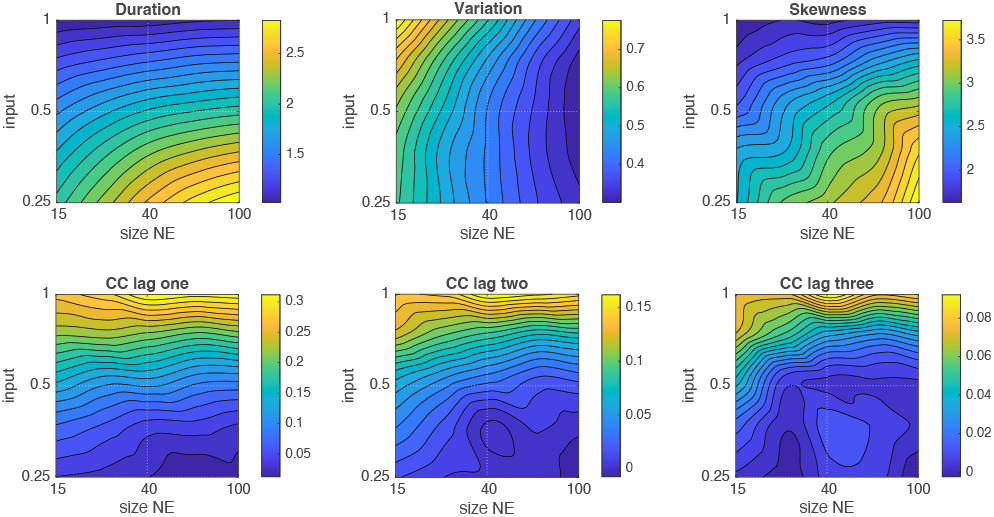
Perception constrains the size of lower-level populations. The values of six reversal statistics – dominance duration, coefficient of variation, skewness, sequential correlations for lag one, two, and three – change differentially with external input and the size of lower level populations, *N*_*e*_.

## Notes

### Competing Interest Statement

The authors have declared no competing interest.

### Summary of Updates

Introduction updated. Discussion updated. New references 40-43 added and relation to present work explained.

### Funding

- Innovationsfonds OvGU

